# RNASeek: A Cross-Phyla Generative Foundation Model for Multipurpose RNA Modeling and Reinforcement Learning-Based Design

**DOI:** 10.64898/2026.09.24.754173

**Authors:** Shiyuan Chen, Neil R. Fernandes, Wei Vivian Li, Lili Wang, Zhenyu Jia, Joy S. Xiang

## Abstract

RNA plays central roles in regulating information flow and provides a versatile substrate for engineering biological functions. While large language models (LLMs) have transformed natural language processing and protein design, a general framework connecting RNA foundation models to functional sequence design remains limited. Here, we present RNASeek, a 1.6-billion-parameter generative foundation model built on a DeepSeek architecture and trained on a cross-phyla transcriptomic corpus for RNA sequence representation and generation. Natural-language tokens enable flexible conditional prediction and sequence design using a unified backbone. RNASeek captures species-specific transcript features and intron–exon boundaries in a zero-shot setting. We then fine-tune RNASeek to predict ribozyme self-cleavage activity and viral mRNA stability, revealing interpretable sequence features associated with function, including ribozyme loop flexibility and stem stability, as well as AU-rich motifs associated with mRNA stability. We use these functional predictors as reward models and apply Group Relative Policy Optimization (GRPO) to update RNASeek’s generation policy toward sequences with desired properties. GRPO-guided generation produces faster-cleaving ribozymes and stability-enhancing 3′ UTRs while satisfying user-specified IUPAC constraints. Experimentally validated RNASeek-generated ribozymes achieve wild-type levels of activity, while RNASeek-generated 3′ UTR sequences exceed the performance of the training data and benchmarked AI-generated 3′ UTRs. Together, RNASeek establishes a unified pretrain–predict–optimize framework that connects learned RNA function to controllable de novo sequence design and provides a general strategy for engineering regulatory RNAs with desired properties.

## Introduction

Over the past decade, deep learning has transformed regulatory genomics and RNA biology by enabling sequence-based prediction of diverse functional outputs. Early convolutional and hybrid architectures can predict regulatory activity from DNA sequence alone, including enhancer/promoter-associated signals and the effects of noncoding variation^1,2^. This principle extended naturally to RNA, as deep neural networks learn sequence determinants of protein–RNA binding specificity directly from RNA sequence^1–3^, and splice-site models such as SpliceAI^4^ predict splice-junction effects from primary sequence context with high accuracy. More recently, transformer-based architectures that incorporate long genomic context have pushed these capabilities further, linking distal sequence features to gene-expression readouts and underscoring the importance of long-range sequence modeling for regulatory biology^5^.

Building on these architectures, foundation models pretrained on massive genome and transcriptome corpora have begun to learn transferable representations that capture evolutionary and species-specific signals^5–7^. Task-focused sequence models further show that sequence alone can predict aspects of mRNA degradation/half-life and related expression phenotypes^8–11^. Such examples include RNA-FM^12^, RNAErnie^13^, SpliceBERT^14^, RiNALMo^6^, Nucleotide Transformer^6,15^, and HyenaDNA^16^. In parallel, generative and combinatorial optimization approaches have emerged to design mRNA sequences with improved stability and translation by combining secondary structure and codon usage, with direct relevance to therapeutic applications. However, most existing RNA language models are designed primarily for representation learning and downstream prediction, rather than for instruction-conditioned sequence generation guided by quantitative experimental rewards.

Reinforcement learning (RL) is a complementary approach for training and aligning generative models toward explicit objectives. Reinforcement learning from human feedback (RLHF)-style pipelines have demonstrated that reward-driven optimization can steer large language models beyond what can be achieved with supervised training alone^17^. In the RNA domain, RL has been applied to specific RNA design problems such as inverse folding, where reward signals encode structural constraints and target functional criteria^17,18^. However, large-scale applications to the *de novo* synthesis of biologically informed RNA transcripts remain limited. More recent policy-optimization variants, including Group Relative Policy Optimization (GRPO), improve upon standard RLHF by reducing variance and computational overhead during large-model optimization, making reward-driven RNA engineering more practical at scale^17–19^. These advances suggest a path toward RNA generative models that can be directly optimized against biologically meaningful reward signals, reducing reliance on repeated time- and resource-intensive wet-lab experimentation or mutagenesis. At the same time, as RNA engineering increasingly moves toward programmable sequence design, there is a growing need for foundation models that can integrate human-specified objectives, biological annotations, and experimental constraints with sequence-based representations. Combining natural-language-style task commands with RNA modeling provides a flexible interface for specifying diverse design goals without redesigning the tokenizer or model architecture for each new task. This leaves a knowledge gap for regulatory RNA design workflows that connect transcriptome-scale generation to quantitative functional annotation rather than target-structure recovery alone^20,21^.

Effective RL-based alignment of RNA generative models requires dense, quantitative reward signals that map sequence variation to functional outcomes across large and diverse sequence spaces. The regulatory architecture of messenger RNAs (mRNAs) makes them particularly well suited for this purpose. Beyond their protein-coding role, mRNAs encode extensive regulatory information in untranslated regions (UTRs), where sequence motifs, base composition, and higher-order RNA structures collectively influence transcript stability and translation efficiency^9–11,22^. Massively parallel reporter assays (MPRAs) can capture these effects across thousands of sequence variants, providing the quantitative sequence–function relationships needed to train and evaluate reward models. To encompass distinct regulatory landscapes, we consider two complementary systems: structurally constrained hammerhead ribozymes and more diverse viral RNA elements embedded in 3′ UTRs. Hammerhead ribozymes provide a mechanistically interpretable sequence–function landscape in which peripheral loop sequence variation modulates a defined self-cleavage activity^23–27^, whereas viral RNA elements encompass a broader and less structurally prescribed space of sequence-dependent effects on RNA stability and translation^22^. Together, these systems provide complementary test cases for evaluating whether a foundation model can learn quantitative regulatory relationships across sequence spaces of increasing complexity.

We introduce RNASeek, a ∼1.5–1.6B-parameter decoder-only foundation model for regulatory RNA tasks that integrates large-scale, cross-phyla pretraining with reinforcement learning–based sequence optimization. RNASeek uses a unified token vocabulary spanning RNA nucleotide sequences, genomic annotation markup, RNA secondary structure representations, and ∼12,000 natural language tokens, providing a unified representation for biological sequence, structure, annotation, and task specifications. Built on a modern GPT architecture, RNASeek is designed to learn transferable transcriptomic representations while supporting quantitative prediction of regulatory activity and controllable generation of functional RNA sequences. We use this framework to connect transcriptome-scale representation learning with sequence–function prediction and reward-driven RNA design, providing a unified approach for modeling and engineering regulatory RNAs.

## Methods

### Foundation Model Architecture and Input Tokenization

#### BPE Tokenization

The input to RNASeek consists of RNA sequences, structural annotations, and natural-language-style task instructions. To encode this multimodal input, RNASeek uses byte-pair encoding (BPE) to segment long text into smaller token units. Traditional RNA models often accept only nucleotide sequences as input and therefore require task-specific retraining because task specifications cannot be conveyed directly through the input prompt. In contrast, the RNASeek tokenizer employs a hybrid strategy that integrates genetic and natural-language representations. RNA tokens include nucleotide sequences, genomic markup, and predicted RNA structure. Natural-language tokens are used to specify task types alongside RNA sequences, supporting command-conditioned prediction and design and enabling extension to additional RNA tasks through continued training or instruction-aware fine-tuning.

#### Token types

1. Nucleotides: Each nucleotide (A, U, C, G) is treated as a distinct token.
2. Predicted RNA secondary structure: Parentheses “(” and “)” are used inline (e.g., (AGU)) to denote base pairing.
3. Natural language: A vocabulary of ∼12,000 tokens, derived from PubMed/PMC abstracts, captures common English words and individual letters (A–Z) to support natural language instructions.
4. Instructions: Task-prefix commands (e.g., ∼$) are used to specify the desired operation.
5. Genomic Markup and Strand-Invariant Markers: We implemented an extensible genomic markup language to explicitly define functional genomic regions. Transcripts are annotated using the following internal markers:
  ● |u5u|: 5′ untranslated region (UTR)
  ● |u3u|: 3′ untranslated region (UTR)
  ● |s|: start codon
  ● |t|: stop codon
  ● |i|: intronic sequence
  These markers enclose the corresponding biological elements (e.g., |s|AUG|s|ACCTG…|t|UAG|t|). They are designed to be palindromic, ensuring that functional boundaries remain consistent when the string is inverted for potential reverse-complementary inversion that might be useful for future tasks.

All components described above are processed through the BPE tokenizer; to the model, they are represented as tokenized sequences as follows:

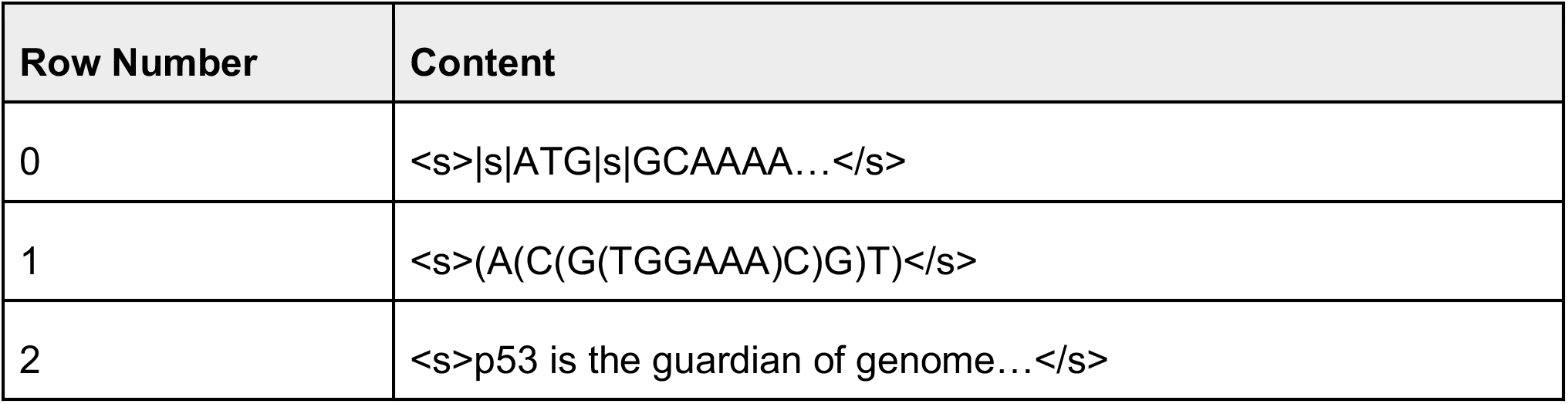

During pretraining, the model predicts the next token at each position.

#### Data Curation and Preprocessing

The pretraining corpus for RNASeek comprises approximately 1.5×10^11^ mRNA tokens (nucleotides) sourced from Ensembl^7,30^. For every genome, the respective GTF file was used to extract transcript regions. The dataset includes mRNA sequences from more than 100 diverse species, spanning the animal, plant, fungal kingdoms, and viruses^31^. This cross-phyla training strategy was designed to capture a broad range of codon biases, regulatory syntaxes, and structural motifs inherent across eukaryotic evolution.

In addition, we incorporated approximately 2 billion natural-language tokens from PubMed abstracts, restricted to entries containing at least six nucleotides within the text body (defined as a contiguous string consisting only of A, U/T, C, or G). This corpus provides textual context for known motifs and encodes natural-language words primarily as lowercase characters, whereas nucleotide symbols are represented in uppercase. This separation helps the tokenizer distinguish biological sequence tokens from natural-language context and supports command-conditioned task specification without redesigning the tokenizer for each new task.

To maximize computational efficiency and capture long-range dependencies, sequences were packed into windows with a context length of 32,768 tokens. To integrate RNA secondary structure information into sequence-level modeling, we augmented approximately 50% of the training sequences with dot-bracket notation predicted using RNAfold^32^ (ViennaRNA version 2.7.0). For brevity, we omitted dots representing unpaired bases and retained only brackets to signal interactions. For example, a sequence and its predicted fold would be interleaved as follows: A(U(GG)C)A. In this notation, nested brackets denote specific Watson-Crick or wobble base-pairing interactions. By linearizing structural information in this manner, the transformer self-attention mechanism can learn dependencies between primary nucleotide composition and predicted secondary structure; this structural encoding was used for both ribozyme and stability prediction.

#### Model Configuration

RNASeek uses a GPT-style decoder-only transformer architecture. Unlike encoder-style models, which use bidirectional self-attention to condition on the full input sequence, decoder-only models use causal self-attention, allowing each token to attend only to preceding tokens during next-token prediction. The RNASeek transformer backbone was initialized from a DeepSeek 1.2B model, which follows the Qwen2.5 architectural design and contains 28 transformer layers^33^.

### Pretraining (Causal Language Modeling)

During pretraining, the model was optimized using standard Next-Token Prediction (NTP) with a Cross-Entropy loss:

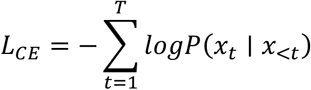

where *ℒ*CE denotes the cross-entropy loss over the sequence, T is the total length of the tokenized input sequence, and t indexes token position. We define x*₀* as the beginning-of-sequence token <S>, so that x<t denotes the context available before position t, i.e., x*₀*, x*₁*, …, xₜ*₋₁*. Thus, for the first sequence token, P(x*₁* | x<1) corresponds to P(x*₁* | <S>). The term P(xₜ | x<t) is the conditional probability assigned by the model to token xₜ given its preceding context. The objective minimizes the negative log-likelihood of the observed sequence, encouraging the model to assign high probability to the correct next token at each position. A 9:1 training:validation split was used on the total corpus.

#### Hardware and Hyperparameters

To ensure the reproducibility of our training pipeline and characterize the computational framework required for large-scale RNA modeling, we have detailed our hardware configuration and optimization parameters below. These settings were selected to balance high throughput with the intensive memory requirements of the 32,768 token context window. The following table summarizes the infrastructure, optimization regime, and parallelism strategies employed to stabilize the 1.6B parameter backbone during both the pretraining and reinforcement learning phases.

| Parameter | Value |
| --- | --- |
| Hardware | 8x NVIDIA A100 (80GB) |
| Interconnect | NVLink |
| Precision | bfloat16 |
| Optimizer | AdamW ($\beta_1 = 0.9$, $\beta_2 = 0.95$, $\epsilon = 1e-8$) |
| Learning Rate | $5 \times 10^{-5}$ (with Cosine Decay) |
| Parallelism Strategy | DeepSpeed (ZeRO-3), FSDP, FlashAttention-2 |

### Supervised Fine-Tuning (Regression)

#### Massively parallel reporter assay datasets

Ribozyme library datasets include the c-di-GMP ribozyme library from Xiang et al^23^, where Loop I contains the c-di-GMP aptamer and Loop II is fully randomized. Ribozyme activity was quantified as the ratio of RNA to DNA reads for each library sequence, obtained using RNA/DNA-seq. Reporter expression data of 57,560 ribozyme sequences with both Loop I and Loop II randomized were generated via FACS-seq^24^. Briefly, cells harboring individual ribozyme sequences cloned downstream of a GFP reporter were sorted into six activity bins based on GFP/mCherry ratios by FACS. Sequencing reads for each ribozyme sequence from each bin were counted, and ribozyme activity was estimated using a lognormal distribution model. The dataset of RNA stability measurements for viral RNA fragments was obtained from Seo et al^22^. These RNA sequences are inserted in the 3′ UTR and their effect on RNA stability was quantified using the same metric as in the c-di-GMP ribozyme dataset, i.e. the RNA/DNA read count ratio.

For ribozyme efficiency and 3′ UTR stability prediction, target values (y) were transformed using a log-plus-one z-score:

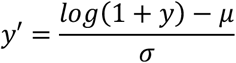

where y denotes the original target value (e.g., measured ribozyme efficiency or 3′ UTR stability), and log(1 + y) is applied to reduce skewness and stabilize variance, particularly for non-negative and heavy-tailed distributions. The transformed value is then standardized using the dataset statistics, where μ is the mean and σ is the standard deviation of log(1 + y) computed over the training set. The resulting y’ represents the normalized target used for regression, ensuring approximately zero mean and unit variance to facilitate stable and efficient model training. As with the pretraining, a 9:1 training:validation split was used.

#### Train–validation sequence-space overlap control

For the ribozyme benchmark, similarity between the training and validation splits was assessed using only the variable regions so that the three invariant catalytic-core segments did not inflate sequence identity. The core sequences GCTGTCACCGG, TCCGGTCTGATGAGTCC, and GGACGAAACAGC were removed, leaving Loop I and Loop II for comparison. Each of the 5,756 validation examples was compared exhaustively with all 51,804 training examples; Levenshtein distances were calculated separately for the two loops, summed, and normalized by their combined lengths to obtain nearest-neighbor sequence identity. No validation example had an exact match to a training example across both variable regions. Normalized 5-mer frequency profiles from Loop I and Loop II were also analyzed by PCA, with PCA fitted on the training set and validation examples projected into the fitted space (Supplementary Figure 1).

For the viral 3′ UTR stability benchmark, constant cloning flanks and the 7-nt barcode were removed before similarity filtering, leaving the 130-nt viral sequence. To reduce train–validation leakage, MMseqs2 was used for forward-strand local nucleotide alignment with the Smith–Waterman algorithm to compare each original training sequence with the validation set. A training sequence was excluded if an alignment to any validation sequence satisfied both at least 50% nucleotide identity and at least 50% query and target coverage. This removed 4,669 of 23,484 original training sequences, leaving 18,815 training sequences and 2,935 validation sequences. As an orthogonal design-space check, normalized 5-mer frequency embeddings were analyzed by PCA using the retained training set as the fitted basis (Supplementary Figure 2).

#### Model architecture

A single-layer feed-forward network (FFN) regression head was appended to the pooled hidden state of the final layer (layer 28) of the RNASeek backbone. The model was trained to minimize the mean squared error (MSE) loss:

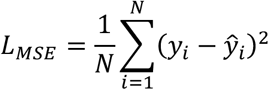

where *ℒ*MSE denotes the mean squared error loss over a batch of samples, N is the total number of samples in the batch, and i indexes each individual sample. yᵢ represents the ground-truth target value for sample i (after any preprocessing such as normalization), while ŷᵢ is the corresponding prediction produced by the model’s regression head. This objective optimizes continuous-valued regression prediction by penalizing the squared deviation between prediction and ground-truth target values.

#### Fine-tuning Data Format

Each fine-tuning example consists of a tokenized RNA sequence paired with a task instruction token that specifies the regression target. Two representative examples are shown below:

*Ribozyme efficiency prediction:*

<S>AACTGG…TTGAAG∼$predict_efficiency\n</S>

*Stability prediction:*

<S>AACTGGAT… TTTGAAG∼$predict_stability\n</S>

In both cases, the regression target (stability or ribozyme efficiency) is derived from the pooled hidden state of the final layer after the</S> token position (EOS pool). The task instruction token (e.g., $predict_stability, $predict_efficiency) conditions the model on the desired output, enabling a single model to perform multiple regression tasks within a unified framework.

#### Baseline models for predicting ribozyme cleavage efficiency and stability

One of the baseline models was a two-layer Long Short-Term Memory (LSTM) network. This model used a simple one-hot encoding for the four nucleotides and three dot-bracket symbols, i.e., “(”, “.”, and “)”, denoting RNA secondary structure. The first LSTM layer accepts an input of 113 features and projected them to a hidden size of 256, followed by a second LSTM layer that expanded the representation to 512 hidden units. The output of the second LSTM was passed through a ReLU activation and a dropout layer (p = 0.2) for regularization. The resulting tensor was flattened and passed through two fully connected layers (3,584 → 1,000 → 1) to produce a scalar prediction. The model was trained using the Adam optimizer with mean squared error (MSE) loss; the learning rate (4 × 10*⁻*⁶) was determined empirically.

Additional baseline models were pretrained DNA/RNA models selected for relevance to UTR-related tasks and feasibility within an academic computing environment. Each model was trained using the standard Hugging Face Trainer with a regression head and author-recommended settings when available. When recommended settings were unavailable, we used a batch size of 16 and a learning rate of 5 × 10*⁻*⁵, which are the same parameters used in RNASeek regression modeling.

All benchmark models, including RNASeek, were trained for a maximum of 200 epochs with early stopping (patience = 20), meaning that benchmark training was terminated when validation performance failed to improve for 20 consecutive epochs. This limit was selected as a conservative upper bound rather than an expected training duration, as pretrained transformer models are commonly adapted to downstream tasks using substantially shorter fine-tuning schedules. For example, the original BERT study evaluated fine-tuning over 2–4 epochs^34^, while downstream fine-tuning schedules reported for the RNA language model RiNALMo ranged from 2 to 50 epochs depending on the task^6^. Ribozyme benchmark performance was summarized by Pearson correlation on the validation set. For the stability benchmark, Spearman rank correlation was calculated at the individual-sequence level across the complete validation set because the measured stability distribution contained a pronounced central mass

### Reinforcement learning for generative RNA design

#### Supervised adaptation for directive guided sequence design: Continued pretraining and IUPAC instruction-aware training

For the 3′ UTR reinforcement learning design task, we performed continued pretraining of RNASeek using human 3′ UTR sequences extracted from the Ensembl hg38 annotation. This step shifted the model generation distribution toward human 3′ UTR-like sequence grammar. We next trained the model to follow IUPAC-based sequence constraints by pairing random nucleotide sequences with computationally generated IUPAC inclusion or exclusion instructions for 50,000 steps. This instruction-aware stage was designed to teach the model how to satisfy computationally specified motif-level constraints rather than simply memorizing naturally occurring RNA patterns from pretraining.

The ribozyme-generation task used a different adaptation strategy because the target design space is defined by compact ribozyme architecture rather than human 3′ UTR grammar. Therefore, ribozyme generation did not include continued human-genome pretraining. Instead, the model was trained directly on computationally generated random sequences paired with matched IUPAC constraints, and subsequent optimization was guided by the RNASeek ribozyme regression model.

#### Reinforcement Learning Policy

To enable reliable generation of stable 3′ UTR and ribozyme sequences, we guided the pretrained RNASeek language model using Group Relative Policy Optimization (GRPO), with the RNASeek regression model serving as the reward function. Each training prompt began with a beginning-of-sequence (BOS) token |u3u|, which also served as an end-of-sequence (EOS) marker to signal completion of generation. Each prompt additionally included a natural-language task instruction of the form ∼$predict_3putr_… or ∼$predict_ribozyme_…, conditioning the model on IUPAC seeds that specify sequences to include or exclude during generation (e.g., restriction sites for cloning, miRNA seed sequences, or RNA-binding motifs). Representative prompt formats are shown below:

<S>∼$predict_utr_exclude_ACT_include_GCT\n

After the newline character (\n), the language modeling head completes the predicted stable UTR region or candidate ribozyme sequence. The EOS token</S> is absent because generation is terminated by the |u3u| marker.

<S>∼$predict_ribozyme_exclude_ACT_include_GCT\n

After the newline character (\n), the language modeling head completes the predicted ribozyme sequence. The EOS token</S> is similarly absent; otherwise, the decoder interprets the input as complete and terminates generation immediately.

#### Reinforcement learning using GRPO

At each GRPO step (Figure 5A), the policy generated 16 candidate sequences using sampling temperature 1.2, top-p = 0.9, and a maximum rollout length of 250 tokens. Each generated sequence was scored using a composite reward consisting of (i) the RNASeek regression-model score and (ii) a syntax/structure score. The regression model was used as the reward model (RM) for both ribozyme activity and 3′ UTR stability optimization. The syntax/structure score reflects adherence to the BOS/EOS |u3u| markers, fidelity to IUPAC constraints, nucleotide composition validity, and a target length of 137 nt. Because the regression training libraries were centered near 137 nt, the length component encouraged generated sequences to remain within the reward model’s best-supported sequence-length regime. The final scalar reward combined 0.4 × RM score and 0.6 × syntax/structure score, prioritizing valid and constraint-compliant generations before reward-model optimization. These rewards were used to update the policy for 75,000 GRPO steps at a batch size of 16.

#### Data, natural language prompting, and rewards

The reward signal was a linear composite of the following components:

1. Length score (targets pure nucleotide sequence length of 137 nt; maximum rollout length = 250 nt; rollouts exceeding 250 nt were discarded):

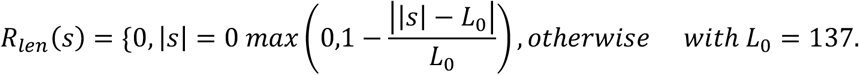
2. Noise/grammar score (penalizes non-ACGT characters in the raw extracted body string):

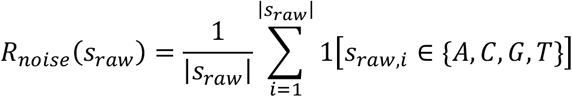
3. Marker score (boundary marker correctness):

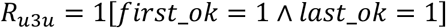
4. Motif inclusion signal is calculated as:

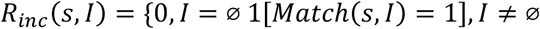 Fraction matched refers to the proportion of generated sequences that can be resolved to the IUPAC motif in their prompt according to the above metrics.
5. Motif exclusion signal is calculated as:

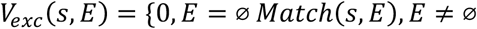

#### Ablation and control-sequence generation

To evaluate whether RNASeek-generated sequences depended on learned sequence organization rather than simple base composition, we generated composition-matched shuffled controls. For each RNASeek-generated sequence, nucleotide order was permuted while preserving the original nucleotide composition; for the ribozyme analysis, loop length and GC content were additionally matched where specified. Throughout the manuscript, “shuffled” refers exclusively to controls derived by permuting an existing sequence, whereas “random” refers to independently generated random nucleotide sequences. Comparisons between RNASeek-generated sequences and their shuffled controls therefore test whether predicted or measured activity depends on sequence arrangement, motif placement, and higher-order RNA syntax rather than composition alone.

We also included sequences generated by a general-purpose language model as an external baseline. ChatGPT-generated 3′ UTR sequences were collected using the following prompt:

Generate exactly **50 unique RNA sequences intended to function as 3′ untranslated regions (3′ UTRs)** and to maximize transcript stability based on your general biological knowledge of RNA stability, degradation, RNA-binding motifs, nucleotide composition, and secondary-structure tendencies.

These must be **3′ UTR sequences only**. Do not generate 5′ UTRs, coding sequences, promoters, amino-acid sequences, explanations, or annotations inside the sequences.

Requirements:

● Generate exactly 50 sequences.
● Each sequence must be between **150 and 200 nucleotides** long.
● Use only the RNA nucleotides **A, U, C, and G**.
● Make every sequence unique.
● Do not include a terminal poly(A) tail consisting of a long artificial run of A nucleotides.
● Avoid obvious repeated templates or sequences that differ only by a few substitutions.
● Design each sequence independently to be as favorable for transcript stability as possible.

Return the results in FASTA format using headers >3UTR_01 through >3UTR_50. Put each complete sequence on a single line immediately below its header. Before responding, verify that there are exactly 50 sequences, that every sequence is 150–200 nucleotides long, and that every sequence contains only A, U, C, and G. Output only the FASTA records and no additional commentary.

These sequences were used to compare RNASeek’s reward-optimized generation against a non-specialized language-model baseline.

### RL Result Motif Analysis

Motif enrichment analysis was performed using MEME Suite^35^ (STREME) in a discriminative setting, with RNASeek-generated sequences as the foreground and GC-matched human 3′UTR sequences as the background. Prior to analysis, only known technical scaffold and adapter regions were removed, while all other sequence content, including viral-like or repetitive elements, was retained to avoid discarding potentially learned biological features. Human background sequences were greedily matched to the GC-content distribution of the RNASeek foreground to minimize compositional bias.

STREME was used to identify motifs significantly enriched in the RNASeek sequence set relative to the matched background, replacing direct k-mer counting approaches with de novo probabilistic motif discovery. The resulting position probability matrices were parsed and visualized as sequence logos, enabling characterization of sequence patterns preferentially generated by the model while controlling for global nucleotide composition.

### Dual luciferase plasmid construction and activity assays

Plasmids for reporter assays were derived from a dual luciferase reporter system based on ref^36,37^ (Addgene #181934). It encodes *Cypridinia* and *Gaussia* luciferases (CLuc and GLuc) under the control of bidirectional EF1a and CMV promoters, respectively. For ribozyme assays, wild-type sTRSV hammerhead ribozyme and inactive control sequences were designed with each ribozyme flanked by A-rich sequences, as previously described^23^. DNA sequences were synthesized as custom oligonucleotides (ThermoFisher), assembled by PCR, and cloned into the XbaI and XhoI restriction sites of the reporter plasmid. For evaluation of candidate 3′ UTR sequences, each candidate sequence was similarly synthesized as custom oligonucleotides (ThermoFisher), assembled by PCR, and cloned into the XbaI and BamHI restriction sites of the reporter plasmid. The parental reporter plasmid lacking an inserted poly(A) sequence was used as a negative control.

Luciferase assays were performed via transfection of dual reporter plasmids using Lipofectamine 2000 reagent (Invitrogen 11668019) and according to the manufacturer’s protocol. Luciferase activity assays were performed using the UltraBrite Cypridina-Gaussia dual luciferase assay reagent from Targeting Systems (DLAR-4 SG-1000) and measured on a Molecular Devices SpectraMax® iD5 Multi-Mode Microplate Reader using an integration time of 2 seconds/well.

## Results

### Construction of RNASeek as a DeepSeek-based RNA modeling framework

We developed RNASeek, a decoder-only language model with 1.6 billion parameters, built upon a Qwen-distilled DeepSeek backbone and trained with a 32,000-token context window (Figure 1A). The RNASeek tokenizer comprises four token categories. The first category contains single nucleotide tokens, A, C, G, and T/U. These tokens remain one nucleotide long, ensuring single-nucleotide resolution at the input level. The second category contains dot-bracket annotation tokens encoding predicted RNA secondary structure using RNAfold^32^; these are reserved tokens and will not merge with any other token type. The third category contains annotation features, including untranslated region (UTR) markers such as “|u3u|” and “|u5u|”, start/stop codon markers such as “|s|” and “|t|”, and “|i|” for intron markers. The fourth category consists of natural-language command prompts, marked by “∼$”, that specify downstream tasks. This component consists of 12,000 vocabulary tokens drawn from natural language, which enable the model to incorporate new instructions without repeating the resource-intensive pretraining step. Together, this tokenization scheme encodes not only raw nucleotide sequence and secondary structure, but also structural annotations and natural language commands, enabling a unified model capable of multitask optimization across diverse downstream objectives.

**Figure 1.**
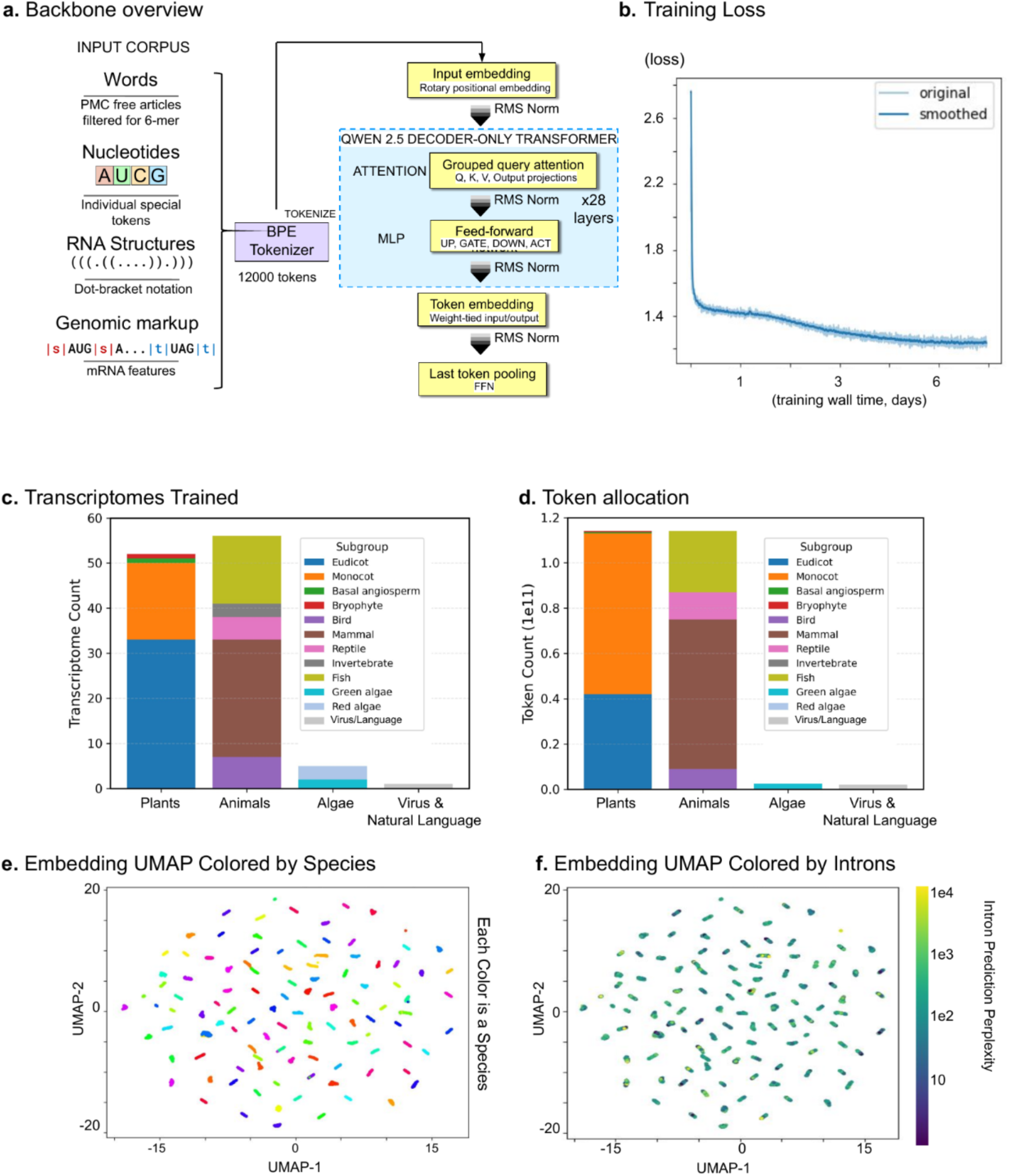
Large-scale pretraining establishes RNASeek as a general RNA foundation model. a. RNASeek architecture, using AUCG nucleotides, natural language tokens, dot-bracket () notation, and genomic markup language tokens as input. Constructed from the DeepSeek 1.5B architecture, which follows a Qwen design. b. Next-token prediction cross-entropy loss during RNASeek pretraining. c. Number of transcriptomes included in RNASeek pretraining. d. Number of transcriptome nucleotides processed during pretraining. Each nucleotide was treated as a token. e. UMAP of next-token prediction embeddings per transcript in each transcriptome, colored by species. Each point represents a transcript. f. UMAP of next-token prediction embeddings per transcript, colored by intron prediction perplexity. Perplexity is defined as the log cross-entropy loss for intron boundary marker prediction.

To ensure the model generalizes across a wide range of RNA motifs, we constructed a cross-phyla training corpus spanning plants, animals, algae, viruses, and natural language (Figure 1C–D). Token allocation was weighted toward better-characterized model organisms while maintaining broad phylogenetic coverage (Figure 1D). Approximately 50% of the overall training budget was assigned to plant transcriptomes, and the majority of the remainder was allocated to animals. In contrast, tokens from algae, viruses, and natural-language text were comparatively scarce. These categories are underrepresented in our model, accounting for only ∼0.5% of the total token allocation and are not expected to drive overall optimization. Nevertheless, they serve an important role by enabling prompt-based control through custom commands without requiring tokenizer retraining or embedding layer resizing.

We trained the RNASeek foundation model on the tokenized transcriptome and natural-language corpus using a 28-layer DeepSeek-R1 transformer backbone with rotary position embeddings, multi-head attention, and gated MLP layers. The model was trained on 8×A100 GPUs over one week, and the loss curve remained stable (Figure 1B). The cross-entropy training loss decreased from 2.6 to 1.3 in the first few days and continued to decrease to 1.2 by the end of training, indicating that performance remained at least partly compute-bound rather than limited solely by data availability or model capacity. Increasing computational resources may therefore further reduce perplexity and improve performance on downstream tasks.

### Emergent embeddings show clear species separation and zero-shot intron prediction

Having established stable optimization and convergence of the foundation model during pretraining, we next asked whether these learned representations encode biologically meaningful sequence information without any task-specific fine-tuning. To test whether the model learns species-specific sequence features without any post-training, we analyzed its internal representations. Using the original validation dataset used in pretraining, which is a 10% held-out dataset, we sampled ∼256 sequences from each of ∼100 species and extracted the last-layer, last-token embeddings generated during the next-token prediction task.

To visualize the relationship among embeddings, we applied UMAP for dimensionality reduction. The projected representations formed compact, well-separated clusters (Figure 1E). Sequences grouped according to their organism of origin, indicating that the pretrained model intrinsically captures species-specific RNA sequence syntax from exposure during pretraining. The tight but distinctive clusters suggest that the model formed a consistent manifold within and across species. Some sequences from one species projected near another species’ cluster, which may reflect evolutionary conservation of these transcripts across multiple organisms.

To quantify model consistency across species beyond evaluation loss on the held-out dataset, we compared RNASeek’s next-token predictions with those of an untrained uniform token selector over the 12,000-token vocabulary. We assessed the extent to which RNASeek captures species-specific sequence features by probing intron prediction. We chose this task because it requires no additional fine-tuning and can be evaluated through causal next-token prediction alone. We evaluated the intron-token prediction perplexity for every species and observed no systematic degradation for any particular species. Overall, the mean perplexity was 323.649 (Figure 1F). Since perplexity (PPL) is related to the average negative log-likelihood (average cross-entropy, H) by *PPL* = *exp*(*H*), where H is the cross-entropy for a given input and PPL is the perplexity. The corresponding geometric-mean probability assigned to the correct next token is thus

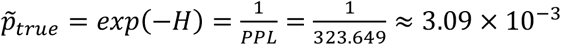

We therefore report an average next-token “recall” (i.e., probability mass on the ground-truth token) of *∼*0.003. This represents a *∼*37-fold improvement over an untrained token selector under a uniform V=12,000 vocabulary, for which

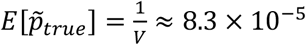

where V is the vocabulary size (12,000) and E is the probability for the chosen token.

We conclude that RNASeek consistently captures sequence structure both within and across species, using intron position prediction as a proxy for cross-species generalization.

### Deciphering Ribozyme Catalytic Efficiency

Building on the stable optimization and broad sequence representations learned during RNASeek pretraining, we next evaluated whether the model could capture sequence–function relationships in regulatory RNA. As a first challenge, we quantified how RNA sequence variation governs ribozyme catalytic activity. Hammerhead ribozymes are RNA-based enzymes that undergo self-cleavage, and this cleavage activity depends critically on tertiary loop–loop interactions between Stem Loop I and Stem Loop II^27,29^. Systematic modulation of these tertiary interactions provides a potential means to quantitatively regulate ribozyme cleavage kinetics. However, modeling RNA tertiary structure remains challenging. We therefore used massively parallel reporter assays (MPRAs) to obtain quantitative measurements of ribozyme activity. Previous MPRAs have mapped tens of thousands to hundreds of thousands of random loop sequences to their self-cleavage activities^23,24^. By inserting these ribozymes into the 3′ UTR of reporter genes, cleavage events reduce reporter expression, allowing ribozyme cleavage efficiency to be inferred from reporter expression levels. These sequence-function datasets provided training data for RNASeek regression models to learn features associated with ribozyme activity.

To establish a performance baseline, we trained a LSTM model and compared it to fine-tuned RNASeek on the task of predicting ribozyme self-cleavage efficiency. The hammerhead libraries were designed such that Stem Loop I and Stem Loop II sequences varied independently, yielding combinations of short and long loop variants (small–small, small–large, large–small, and large–large).

We used the small LSTM model to survey task complexity because it could be optimized quickly, trained rapidly, and did not depend on pretraining. It was evaluated on the same dataset as the final RNASeek model, with target values defined as log1p-transformed z-scores derived from normalized fluorescence measurements from MPRA datasets^23,24^. As ribozyme loop lengths increased from short N4–7 loops to longer N30–60 loops, predictive performance declined (R² = 0.60 → 0.45 → 0.35) (Figure 2A). Longer loop regions expand the combinatorial space of possible base-pairing interactions and tertiary folding configurations, making the sequence–activity relationship more difficult to learn from limited data. These results highlight the limitations of low-capacity sequence models in capturing long-range RNA interactions and motivate the use of higher-capacity architectures with pretrained sequence representations.

**Figure 2.**
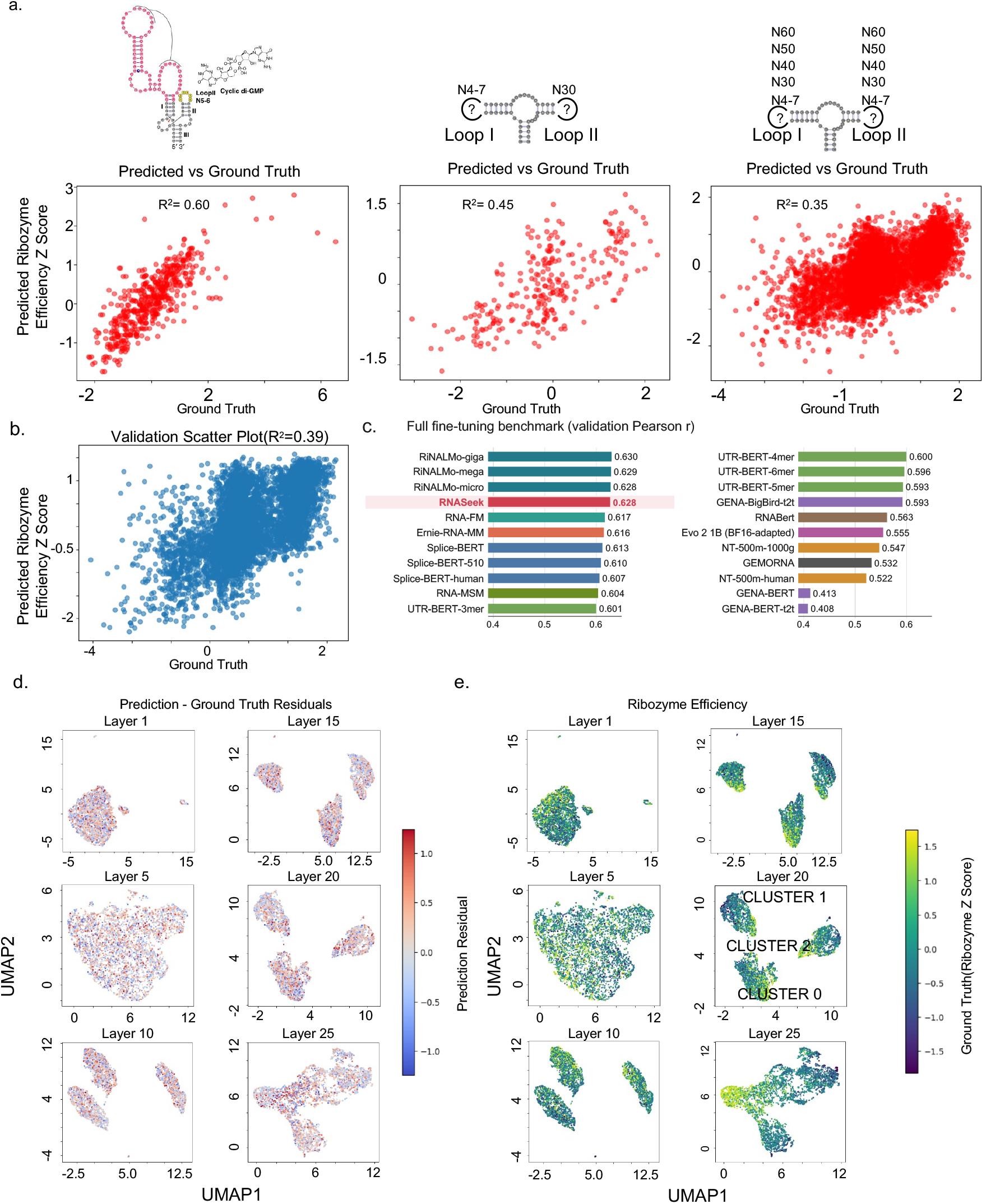
RNASeek predicts ribozyme catalytic efficiency. a. Ribozyme prediction performance compared with the baseline LSTM. For increasingly complex ribozymes, prediction performance is plotted, with the y-axis representing the predicted value and the x-axis representing the ground truth value. Cyclic di-GMP aptazyme activity (left) is given by normalized RNA/DNA levels^23^. Randomized Loop I/Loop II ribozyme activity (middle, right) is given by normalized reporter fluorescence levels^24^. Lower z-score values indicate higher ribozyme efficiency. b. RNASeek prediction performance on ribozyme libraries with two variable loop regions and three constant catalytic core regions. Scatter plot showing predicted (Y) against ground truth (X) values. c. Bar plots comparing RNASeek, GEMORNA, and 18 RNA foundation models on the same task. Each pretrained model was fully fine-tuned for up to 200 epochs with an early-stopping patience of 20, using the model developers’ recommended hyperparameters. Bars show validation Pearson correlation; higher values indicate better performance. d. UMAP of the prediction embeddings for layers 1, 5, 10, 15, 20, and 25, pooled from the last token and colored by prediction residual. A darker color represents a higher residual. e. UMAP of the prediction embeddings for layers 1, 5, 10, 15, 20, and 25, pooled from the last token and colored by ground truth. A darker color represents greater ribozyme catalytic efficiency.

### RNASeek achieves performance competitive with leading RNA foundation models

Compared with the LSTM baseline on the ribozyme efficiency prediction task, the RNASeek regression model achieved higher predictive performance (Figure 2B). We then benchmarked the fine-tuning performance against 19 other foundation models and RNASeek ranked near the top of the benchmarked models (Figure 2C). RNASeek reached a validation Pearson r of 0.628, which is below the BERT-style RiNALMo models, while outperforming the remaining evaluated models. The relatively simple LSTM baseline still ranked above several larger pretrained models, indicating that task-specific inductive bias can remain competitive even when model scale and pretraining differ substantially.

RNASeek uses the same sequence and secondary-structure information as the LSTM baseline but processes these inputs through a decoder-only transformer backbone with natural-language task prompting and last-token pooling. Its benchmark performance therefore demonstrates that causal representations are effective for quantitative ribozyme prediction. More broadly, bidirectional encoders are not strictly required for sequence-level prediction, as causal/autoregressive genomic models such as HyenaDNA have also produced useful representations for downstream predictive tasks.^16^ The decoder-only architecture also provides an important practical advantage for RNASeek: the same pretrained backbone used for prediction can serve directly as the autoregressive policy for prompt-conditioned GRPO sequence generation, preserving the generative capability required for the subsequent design experiments.

We also examined whether model evaluation performance could be explained by train–validation duplication. After removing the three invariant catalytic cores, the 5,756 validation variable regions had no exact matches among the 51,804 training variable regions; the median identity to the nearest training neighbor was 0.658, and 5-mer PCA placed validation examples within the broader training design space without exact overlap (Supplementary Figure 1). Thus, the model was evaluated on closely related but nonidentical ribozyme designs rather than repeated examples.

### RNASeek representations reveal distinct functional submanifolds and sequence features

To understand how RNASeek represents ribozyme sequences and the source of prediction errors, we performed a per-layer embedding analysis. For each layer, we extracted the last-token embedding for every test sequence and applied UMAP to project the representations, coloring each point by its residual (predicted minus ground-truth log1p-transformed z-score; scale held constant across layers). Across early (layers 1–5), mid (layers 10–15), and late (layers 20–25) layers, residuals were evenly distributed across the projected manifold, with no localized regions of consistently large positive or negative error (Figure 2D). Thus, no single layer appeared to introduce a bottleneck that concentrated prediction errors within particular sequence clusters. This observation suggests that RNASeek’s residual errors may partly reflect uncertainty in the underlying sequence–ribozyme activity measurements, while informative representations are progressively and consistently captured throughout the network.

Beyond the distribution of prediction errors, the RNASeek embeddings revealed distinct organization within the ribozyme sequence space. At layer 20, UMAP-based DBSCAN identified three stable clusters spanning sequences with varying cleavage efficiencies (z-scores; Figure 2E). Rather than representing sequences as a single smooth continuum, RNASeek therefore organizes the sequence space into distinct submanifolds that may reflect different mechanistic modes of ribozyme activity. Clustering remained stable across several intermediate layers and became smoother in the final layers, consistent with the integration of discrete sequence representations into the continuous objective of the regression task. These results suggest that RNASeek internally organizes ribozyme sequences into distinct functional groups while retaining the ability to evaluate activity across multiple mechanistic modes.

We next examined sequence and structural features enriched within these clusters to identify potential mechanisms underlying their different activity profiles. We focused on clusters 0 and 2, which contained the broadest ranges of cleavage efficiencies at layer 20. Within each cluster, we compared the top 20% most efficient sequences with the bottom 20% (Figure 3A). A hexagonal heatmap revealed the distribution of ribozyme efficiency as a function of Loop I and Loop II length. In cluster 0, where both variable loop regions were short, high-efficiency enzymes were enriched at Loop I lengths of ∼5–9 nt and Loop II lengths of ∼4–8 nt, with efficiency declining at other Loop I II length ratios. In cluster 2, high-efficiency enzymes were associated with asymmetric designs in which Loop I remained short while Loop II was longer (24–30 nt). Together, these observations suggest that short Loop I and Loop II lengths generally support high ribozyme activity, while a subset of sequences can also achieve efficient cleavage with an extended Loop II.

**Figure 3.**
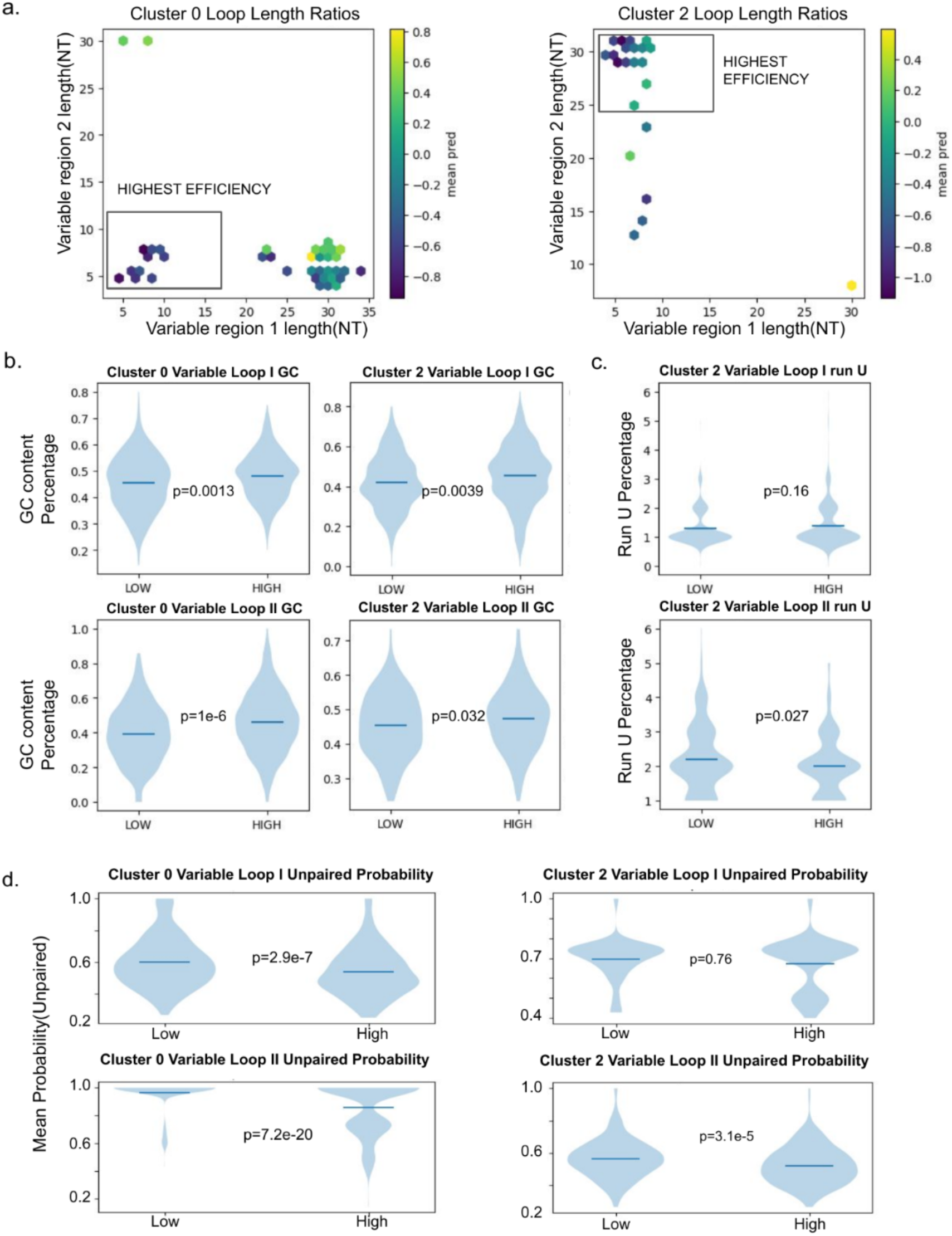
Decoding ribozyme efficiency with RNASeek structural interpretation. a. Mean ribozyme efficiency z-score across all sequence lengths within the same ratio bin; UMAP embeddings were generated from layer 20 and grouped according to their loop length ratio. The x-axis represents the length of loop I, and the y-axis represents the length of loop II. A darker color indicates greater ribozyme catalytic efficiency. b. Violin plot showing the distribution of GC content (number of G/C nucleotides divided by the sequence length) in high-efficiency ribozymes (corresponding to low z-scores); significance values were calculated using Fisher’s exact test. Low – bottom 20% of ribozymes ranked by efficiency (n = 40); high – top 20% (n = 40). c. Poly-U content analysis of high-efficiency ribozymes. The width of each violin represents the local density of observations, and the horizontal line indicates the mean poly-U content. Low – bottom 20% of ribozymes ranked by efficiency (n = 40); high – top 20% (n = 40). d. Violin plot showing the fraction of nucleotides predicted to be unpaired over the sequence length. Base-pairing prediction was performed using ViennaRNA (2.7.0). Low – bottom 20% of ribozymes ranked by efficiency (n = 40); high – top 20% (n = 40).

Subsequently, we asked whether sequence composition and structural accessibility differed between high- and low-efficiency designs. High-efficiency ribozymes consistently exhibited reduced GC content in the variable loop regions (Figure 3B), with significant depletion in both loops in cluster 0 and a weaker but still evident reduction in cluster 2. Consistent with this trend, the high-efficiency group in cluster 2 showed enrichment of U-rich sequences in Loop II, but not Loop I. Contiguous uridines (run-U) are associated with reduced structuredness and greater loop flexibility, and 3-mer run-U sequences were specifically enriched in Loop II of cluster 2 sequences (Figure 3C). To directly test whether cleavage efficiency was associated with reduced base pairing, we used RNAfold (ViennaRNA) to calculate the mean unpaired probability across nucleotides in each loop region. Efficient sequences in cluster 0 showed significantly higher unpaired probability in Loop II, with cluster 2 showing the same trend (Figure 3D). Together with the GC depletion and U-rich sequence enrichment described above, these results indicate that highly active hammerhead ribozymes favor exposed and flexible loops. Importantly, these findings support a permissive design principle in which high cleavage efficiency can be achieved across a broad sequence space without requiring rigid sequence motifs.

### RNASeek generalizes to RNA stability prediction

Although RNASeek demonstrated strong performance in predicting ribozyme activity, ribozyme libraries remain structurally constrained as each variable loop is embedded within constant stem regions. We next examined whether RNASeek could predict the stabilizing effects of viral RNA elements, which span a less constrained sequence space. A recent study^22^ employed an MPRA in which 130-nt RNA fragments tiled across viral genomes were inserted in the 3′ UTR of reporter genes. The abundance of reporter transcripts were then quantified to measure the cis-regulatory effect of each fragment on RNA stability. Using this MPRA dataset, we evaluated whether RNASeek can predict RNA stability as modulated by viral RNA fragments and whether it generalizes to this less constrained sequence space relative to ribozyme activity prediction.

First, we trained a regression head to predict reporter RNA abundance, represented as a log1p-transformed z-score for each MPRA library sequence. After similarity filtering, 18,815 training sequences and 2,935 validation sequences were retained (Supplementary Figure 2). Across the validation set, RNASeek predictions showed a monotonic relationship with measured stability (Spearman r = 0.342; Figure 4A). In the 22-model full-fine-tuning benchmark, RNASeek remained near the top: it matched UTR-BERT-4mer (r = 0.342), which was below RiNALMo-micro (r = 0.348) and UTR-BERT-6mer (r = 0.358), and exceeded the remaining evaluated models (Figure 4B). The advantage of UTR-BERT-6mer may reflect its UTR-specific pretraining and bidirectional 6-mer representation, whereas RNASeek is a cross-phyla autoregressive model designed to preserve general-purpose generative capability rather than maximize specialization for a single UTR prediction task.

**Figure 4.**
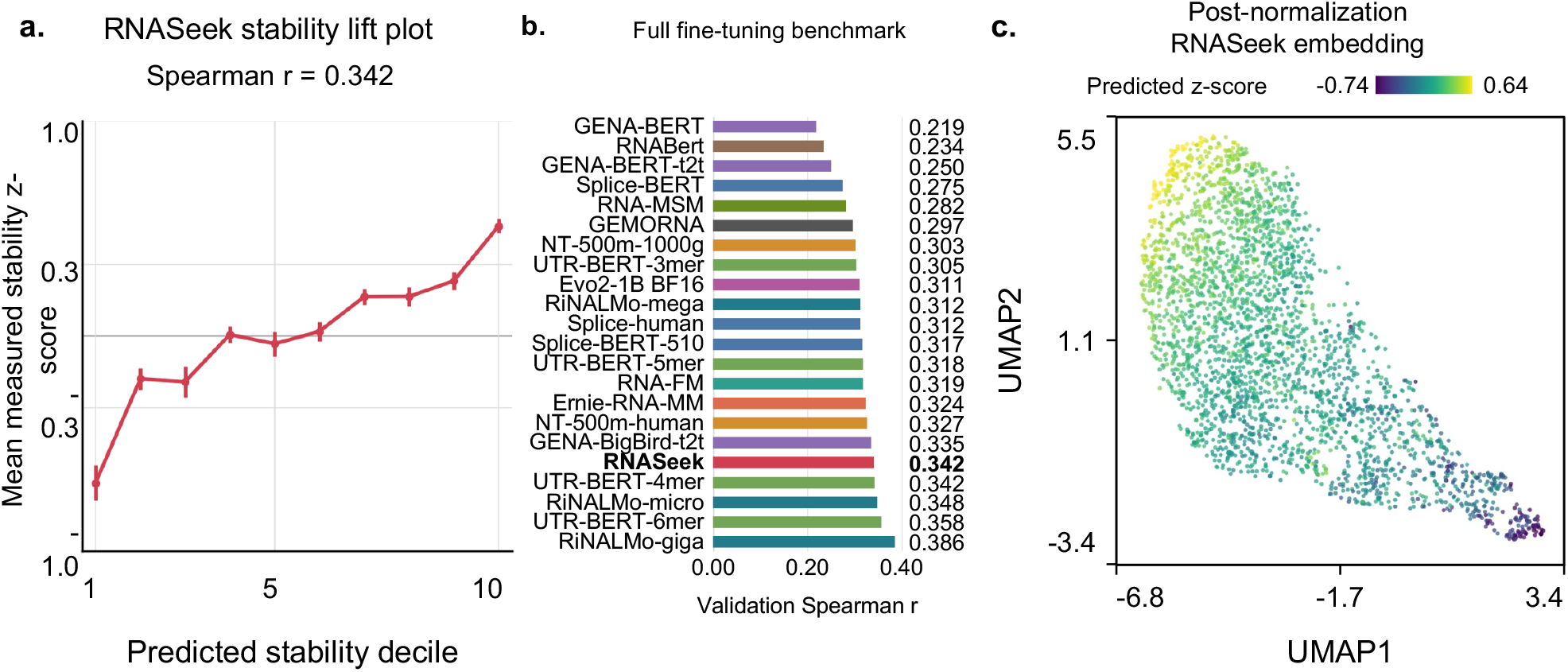
3′ UTR mediated RNA stability prediction with RNASeek. a. RNA stability prediction performance on the dataset from Seo et al. (2023). Predicted stability z-scores were compared with experimentally measured stability across the complete validation set. Sequences were divided into deciles according to predicted stability, and points show the mean experimentally measured stability z-score within each decile with standard error. The Spearman correlation shown in the panel was calculated at the individual-sequence level. Increased predicted stability z-scores are associated with higher ground-truth stability. b. Bar plots comparing RNASeek, GEMORNA, and 18 RNA foundation models on the same stability task. To prevent train–validation leakage, training sequences whose 130-nt variable insert had at least 50% aligned nucleotide identity and at least 50% query and target coverage to any validation insert were excluded. Models were fully fine-tuned for up to 200 epochs with patience 20 using model-specific recommended learning-rate and regularization settings. Bars show Spearman correlation across the complete validation set; higher values indicate better performance. c. UMAP of post-normalization embeddings pooled from the last token of the last layer, colored according to predicted stability z-scores, showing the model’s ranking of sequences in latent space.

Second, to determine whether internal RNASeek representations encode stability-related information, we projected the post-normalization final-layer embeddings before the regression head using UMAP (Figure 4C). Coloring the embedding by predicted stability revealed a smooth gradient across the manifold, indicating that the final transformer representation already organizes sequences according to the stability ranking used by the regression head. Together with the full-set Spearman benchmark, this result supports transfer of the pretrained RNASeek representation to a less structurally constrained regulatory RNA landscape.

### RNASeek enables *de novo* design of stable 3′ UTRs

Having established that the regression framework successfully captures sequence features associated with stability, we next sought to move beyond sequence analysis toward design. Although MPRAs and laboratory-based mutagenesis and evolution can sample portions of sequence–function space, their coverage remains limited. Hence, generating sequences beyond the virome and naturally occurring sequence space could improve our ability to design regulatory elements with desired performance. Such designs could enhance RNA stability for applications in mRNA vaccines, recombinant protein expression, and gene therapy. We therefore used reward-guided RNASeek generation to design stabilizing sequences that may not occur in nature.

To adapt the model for human 3′ UTR design and increase its ability to model human RNA sequence features, we performed continued pretraining of the RNASeek model using 3′ UTR sequences from coding transcripts downloaded from Ensembl (GRCh38). The model was trained for a single epoch without sequence repetition, allowing the model to refine its representation of sequence features and architecture of human 3′ UTRs while preserving the foundational knowledge acquired during initial pretraining.

### GRPO guides RNASeek toward stable 3′ UTRs

For the reinforcement learning (RL) phase (Figure 5A), we used both the generative and regression components of the RNASeek architecture. The generative policy model used the pretrained RNASeek backbone equipped with a language modeling head and sequences were generated stochastically using top-p = 0.9 and temperature = 1.2. Candidate 3′ UTR sequences generated by the policy model were passed to the RNASeek stability regression model, and the resulting predicted stability z-scores were used as reward feedback during Group Relative Policy Optimization (GRPO). This closed-loop design iteratively updated the policy toward generating sequences with higher predicted stability while retaining sequence-generation capability.

**Figure 5.**
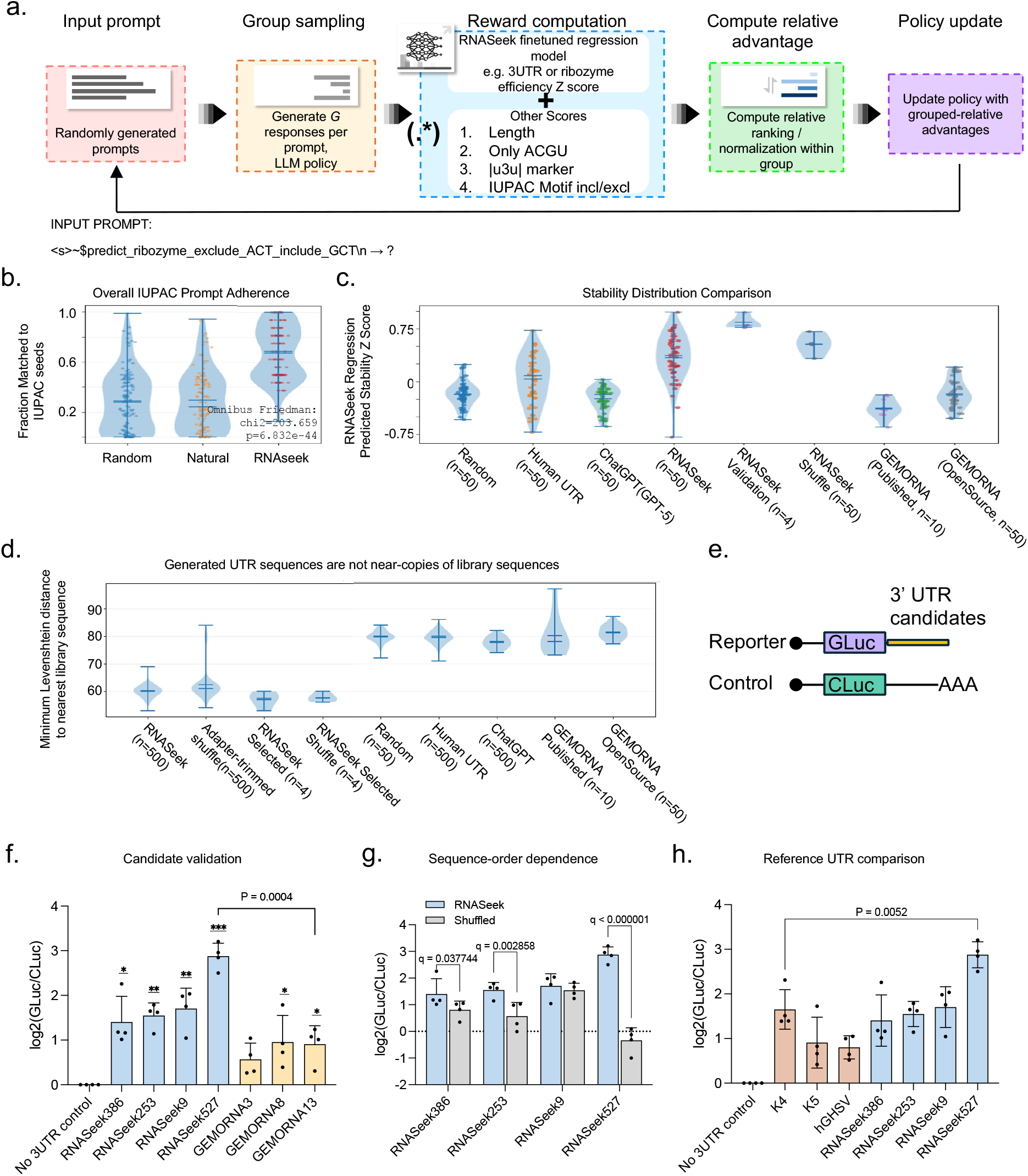
Stable sequence generation with RNASeek. a. GRPO training loop. One loop represents a single training step, during which the model is tasked to complete a randomly generated natural-language prompt by generating a candidate 3′ UTR sequence. The generated sequences are evaluated by the trained regression reward model together with length and prompt adherence using regular-expression matching. For each prompt group, 50 sequences were sampled. b. Post-training evaluation of instruction-following performance measured by sequence motif inclusion and exclusion tasks using IUPAC constraints in the prompt. RNASeek generated sequences were compared against randomly generated sequences and human 3′ UTR sequences. The y-axis indicates prompt adherence as a proportion of sequences satisfying the specified constraints; higher values indicate better performance. For each of the groups compared, 50 sequences were sampled. c. Post-training evaluation of predicted RNA stability, comparing random sequences, natural human 3′ UTRs, ChatGPT-5-generated sequences, RNASeek-generated sequences, the top four RNASeek candidate sequences selected for experimental validation, shuffled RNASeek candidates, published GEMORNA sequences, and sequences generated using the released open-source GEMORNA model. The y-axis indicates predicted stability z-score; higher values indicate better performance. The number of sequences compared in each group is indicated by n. d. For each tested/generated 3′ UTR sequence, the minimum Levenshtein distance to the closest sequence in the regression training set used by regression modeling was computed using the raw full-length sequence. Lower distances indicate greater similarity to at least one training sequence. e. Dual-luciferase reporter assay used to validate candidate 3′ UTR activity. 3′ UTR candidate sequences were inserted downstream of GLuc, and CLuc was used as an internal transfection/expression control. GLuc signal was normalized to CLuc to quantify the effect of each candidate 3′ UTR on reporter expression. f. Experimental validation of RNASeek-designed 3′ UTR candidates. GLuc/CLuc luciferase activities are normalized to the negative control and log*₂*-transformed, showing RNASeek-generated candidates and GEMORNA-derived top ranked sequences. Bars indicate mean±SD. Asterisks indicate two-sided one-sample t-tests against zero on the log*₂*-normalized values (* P < 0.05, ** P < 0.01, *** P < 0.001); the P value for comparing RNASeek527 and GEMORNA13 is given by Welch’s two-sided t-test. g. Comparison of GLuc/CLuc luciferase activity between RNASeek-generated candidates and their corresponding shuffled sequences, which preserved nucleotide composition while disrupting sequence order. Bars indicate mean ± SD of log*₂*-normalized activities. Two-sided unpaired t-tests were corrected across the four comparisons using the two-stage Benjamini–Krieger–Yekutieli false-discovery-rate procedure; significant adjusted values are shown as q values. h. GLuc/CLuc luciferase activities comparing top RNASeek-generated candidates and top sequences from the training data (Seo et al^22^). Bars indicate mean±SD of log*₂*-normalized activities. The P value for RNASeek527 versus K4 is given by Welch’s two-sided t-test.

### GRPO preserves sequence-constraint adherence

To measure how well the trained model followed include/exclude directives, we quantitatively assessed the post-GRPO reliability of RNASeek for motif exclusion and inclusion. For each assessment category, we generated 50 random prompts, each containing one IUPAC inclusion directive and one exclusion directive. We compared three sources of sequences: GRPO-optimized RNASeek, length- and GC-matched randomly generated sequences, and natural human 3′ UTR sequences from Ensembl BioMart. Each generated sequence was evaluated against its corresponding prompt and rated according to inclusion fidelity (at least one appearance according to IUPAC, Hamming distance ≤ 1), exclusion fidelity (IUPAC motif not expressed by the sequence), and overall fidelity (satisfied both metrics at the same time). In the 50-sample test, RNASeek showed higher instruction-following fidelity than random and natural sequences (Figure 5B). The exclusion benchmark also confirmed that RNASeek did not increase excluded-motif occurrence relative to natural sequences. These results indicate that the GRPO pipeline can enrich desired IUPAC motifs while maintaining exclusion performance comparable to random and natural sequences.

### GRPO enriches predicted stability of generated 3′ UTRs

To evaluate whether GRPO improved RNASeek’s ability to design stable 3′ UTRs, we scored generated and control sequences using the previously trained 3′ UTR stability regression model (Figure 5C; Supplementary Data 1). We compared length- and GC-matched random sequences; length- and GC-matched human 3′ UTR sequences from Ensembl; ChatGPT-5-generated sequences; GRPO-optimized RNASeek sequences; four RNASeek sequences selected for experimental validation and their shuffled variants; and GEMORNA sequences. GEMORNA is a decoder-based generative model designed for similar RNA sequence-generation tasks, but is substantially smaller than RNASeek and was developed without modern reinforcement learning approaches such as GRPO. Specifically, we evaluated both published GEMORNA sequences and sequences generated using the released open-source GEMORNA model. Because most existing RNA foundation models are masked-language, BERT-style models and are not naturally suited for de novo sequence generation, we included ChatGPT-5 as a general-purpose GPT-style sequence-generation baseline. This comparison allowed us to test whether a frontier general-purpose language model could generate stable 3′ UTR-like sequences without task-specific RNA optimization.

Across groups, the predicted stability distributions differed strongly. GRPO-optimized RNASeek sequences showed substantially higher predicted stability than random sequences, human 3′ UTR controls, and ChatGPT-5-generated sequences. The four RNASeek sequences selected for validation were among the highest-scoring designs, while their shuffled counterparts showed reduced predicted stability, indicating that the stability signal depends not only on base composition but also sequence order. For GEMORNA, we separately considered the published sequences and sequences generated from the released open-source model. Both the published and open-source model-generated GEMORNA sequences showed lower predicted stability than RNASeek. Together, these results indicate that the blended GRPO objective oriented RNASeek toward stable 3′ UTR design while preserving sequence-level structure beyond simple GC or length effects.

### Generated designs are distinct from training sequences

To test whether RNASeek-generated 3′ UTR sequences were near copies of the regression training set, we computed the minimum Levenshtein distance from each tested sequence to its nearest neighbor in the regression training data used to train the reward model (Figure 5D). RNASeek-generated sequences showed lower nearest-training distances than unrelated random, human 3′ UTR, ChatGPT, and GEMORNA controls, consistent with RNASeek sampling from a sequence space more similar to the training distribution. However, these distances remained large relative to the approximately 137-nt sequence length, and no exact or near-exact matches were observed. Importantly, RNASeek-generated sequences, including the selected candidates, showed distance distributions comparable to matched shuffled controls. This suggests that the observed similarity is better explained by shared nucleotide composition rather than memorization of regression training examples. Together, these results provide evidence that RNASeek does not generate sequences through memorization or near-copying of the regression training set.

### RNASeek-designed 3′ UTRs enhances reporter expression experimentally

To test whether reward-guided RNASeek designs produce measurable activity in cells, we selected four RNASeek-generated 3′ UTR candidates for dual-luciferase reporter validation. In this assay design, each candidate 3′ UTR sequence was cloned downstream of the GLuc reporter, while CLuc served as a transfection internal control for normalization (Figure 5E). Each RNASeek candidate was tested together with a directly derived shuffled sequence that preserved nucleotide composition while disrupting sequence order, four GEMORNA-derived comparison sequences, and negative and positive controls. Reporter activity was measured 48 h after transfection and normalized to the negative-control mean (Figure 5F–H).

All four RNASeek candidates showed greater reporter activity than the negative control, with fold activation ranging from approximately 2.8- to 7.5-fold (Figure 5F). RNASeek527 showed the strongest reporter activity and was significantly higher than the negative control (p=0.0003). We included sequences generated by GEMORNA as a benchmark and found that the RNASeek candidates showed higher activity overall, with RNASeek527 exhibiting significantly greater activity than all tested GEMORNA sequences (p<0.0001; Figure 5F).

Three of the RNASeek candidates showed higher activity than their directly paired shuffled derivatives (Figure 5G). Because each shuffled sequence was derived from the corresponding RNASeek sequence, the decrease after shuffling indicates that the observed activity is not explained by nucleotide composition alone. Instead, the result suggests a contribution from sequence organization, potentially including motif placement, local RNA structure, and higher-order regulatory syntax learned during reward-guided generation.

To test whether RNASeek could design sequences with RNA stability beyond the range represented in the training data, we tested K4 and K5, which were sequences in the library validated to show the highest reporter expression, as well as hGHSV, the top-performing sequence from the complete RNA stability dataset^22^. RNASeek9 matched the expression level of K4, the highest-performing of the tested sequences, whereas RNASeek527 showed significantly greater expression than all tested sequences (p<0.0015). RNASeek527 was also the only sequence that further enhanced reporter expression when followed by the SV40 poly(A) sequence (Supplementary figure 3). These results suggest that RNASeek can generate sequences with reporter expression levels exceeding that of the sequences represented in its training data.

Although RNASeek527 performed particularly well, performance varied across candidates. This variability is expected because reward-optimized sequences are selected using a learned reward model trained on prior sequence-function measurements, whereas reporter activity in a new cellular setting can depend on vector context, transcript processing, RNA-binding protein availability, motif spacing, local secondary structure, and time-dependent reporter dynamics. Thus, RNASeek should be interpreted as enriching the design space for experimentally active 3′ UTR candidates rather than guaranteeing uniformly high stability for every generated sequence.

### RNASeek designs reveal compositional and structural determinants of stability

To evaluate overall sequence composition (Figure 6A), we compared RL-optimized RNASeek sequences against UTR background sequences from Ensembl. RNASeek enriched AU-rich sequences, including motifs resembling AAUAAA polyadenylation recognition sequences, consistent with regulatory features associated with poly(A) tail addition and mRNA stabilization. Notably, while RNASeek outputs were AU-rich, they maintained significantly higher nucleotide diversity than ChatGPT-5 outputs (Figure 6G) and occupied a broader sequence-embedding space. This indicates that reinforcement learning optimized sequence composition for stability while preserving greater sequence complexity than a general-purpose LLM on this specialized RNA task.

**Figure 6.**
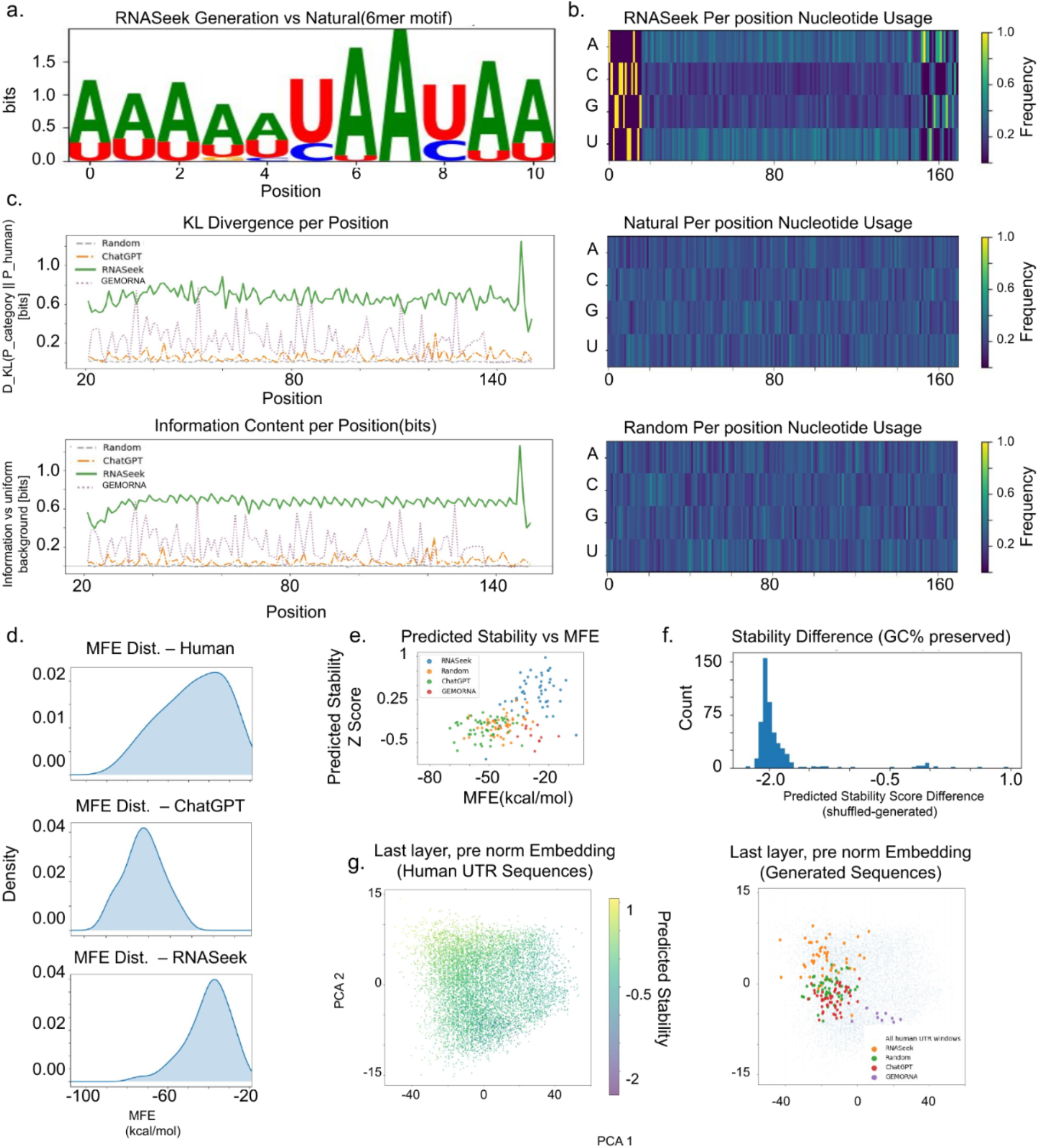
RNASeek stable sequence analysis. a. Sequence logo generated from analysis of RNASeek-generated stabilizing 3′ UTRs. Letter height is proportionate to its frequency in the generated sequences. b. Heatmap showing positional nucleotide frequencies (A, U, C, and G) across RNASeek-generated sequences. Brighter colors indicate higher nucleotide frequency at the corresponding position. c. Positional information entropy relative to random sequences. Higher entropy indicates greater deviation from a random sequence distribution. d. Distribution of minimum free energy (MFE) for each set of sequences. e. Scatter plot showing the relationship between MFE (x-axis) and predicted stability z-scores (y-axis). f. Stability histogram showing the change in predicted stability after sequence shuffle. The x-axis is the stability z-score difference between each shuffled control and its corresponding generated sequence. g. PCA projection of pooled last layer, last-token, pre-head embeddings for natural human 3′ UTR sequences (left), with synthetic sequences projected onto the embedding landscape defined by natural human 3′ UTRs (right).

Positional nucleotide analysis showed that RNASeek-generated sequences exhibit a clear AU-rich bias across the sequence, whereas natural and random controls maintain near-even nucleotide usage apart from minor end-position coverage effects, indicating that RNASeek strategically deploys AU-rich sequence syntax to shift transcripts toward more stable states (Figure 6A–B). Comparison of per-position nucleotide distributions against natural sequences further showed that this bias is not merely inherited from native UTR composition, but instead represents an engineered signal. Kullback–Leibler (KL) divergence analysis showed that many positions in RNASeek-generated sequences carry approximately 0.44 additional bits of information relative to natural controls (Figure 6C).

To determine how this compositional bias relates to RNA structure, we examined the predicted minimum free energy (MFE) of RNA secondary structure folding and found that RNASeek-generated sequences converged on a distinct MFE band rather than the broader distribution observed in natural and random sequences (Figure 6D), suggesting that this folding regime emerged during reinforcement learning rather than continued pretraining alone. Regression analysis (Figure 6E) further showed that MFE was strongly associated with predicted stability up to an apparent ceiling, whereas natural transcripts showed a weaker MFE–stability relationship. These observations are consistent with a model that optimizes folding only to the extent needed to improve predicted stability while preserving sequence diversity and motif compatibility. In silico mutagenesis provided direct support for a role of AU-rich syntax: when AU bases were exchanged with GC while preserving CpG dinucleotide content, stability scores shifted sharply downward, with most perturbed sequences dropping to roughly –2 standard deviations in the reward model (Figure 6F).

Finally, RNASeek-generated sequences occupied a broader region of the latent embedding space than ChatGPT-5-generated sequences when projected alongside random and natural sequences. ChatGPT-5-generated controls clustered within a compact, intermediate-stability region, consistent with a narrower distribution of generated sequences. In contrast, random controls spanned a similarly broad region of the embedding space as RNASeek-generated sequences, indicating that RNASeek retained the ability to generate diverse sequences while shifting them toward higher predicted stability. Together, these findings indicate that the RNASeek reinforcement learning pipeline promotes both predicted stability and sequence diversity without compromising either objective (Figure 6G).

### Reward-guided reinforcement learning generates more active ribozymes

Next, we applied reinforcement learning to the challenge of generating fast-cleaving ribozymes, whose catalytic activity depends on sequence-encoded structural features that are difficult to infer from sequence alone. We coupled RNASeek to the previously trained ribozyme reward model within a closed generator–evaluator loop. In this framework, pretraining provided a broad prior over RNA syntax, while GRPO iteratively amplified generated sequences with features associated with higher predicted catalytic activity. This design parallels the stability RL pipeline, with the generative RNASeek model producing candidate sequences and the regression RNASeek model scoring each candidate for policy optimization.

The generated ribozymes also showed a higher predicted activity distribution than shuffled outputs, IUPAC-resolved sequences, and the original wet-lab training data (Figure 7A; Supplementary Data 2). Because retention time inversely reflects ribozyme activity, shorter retention indicates faster self-cleavage. The loss of predicted activity after shuffling the variable regions in Loops I and II indicates that the performance gain is not explained by simple nucleotide composition alone, but depends on sequence ordering and long-range dependencies captured by the model. Together, these results indicate that RNASeek learned sequence-order and structural features predictive of ribozyme activity rather than simply reproducing training examples.

**Figure 7.**
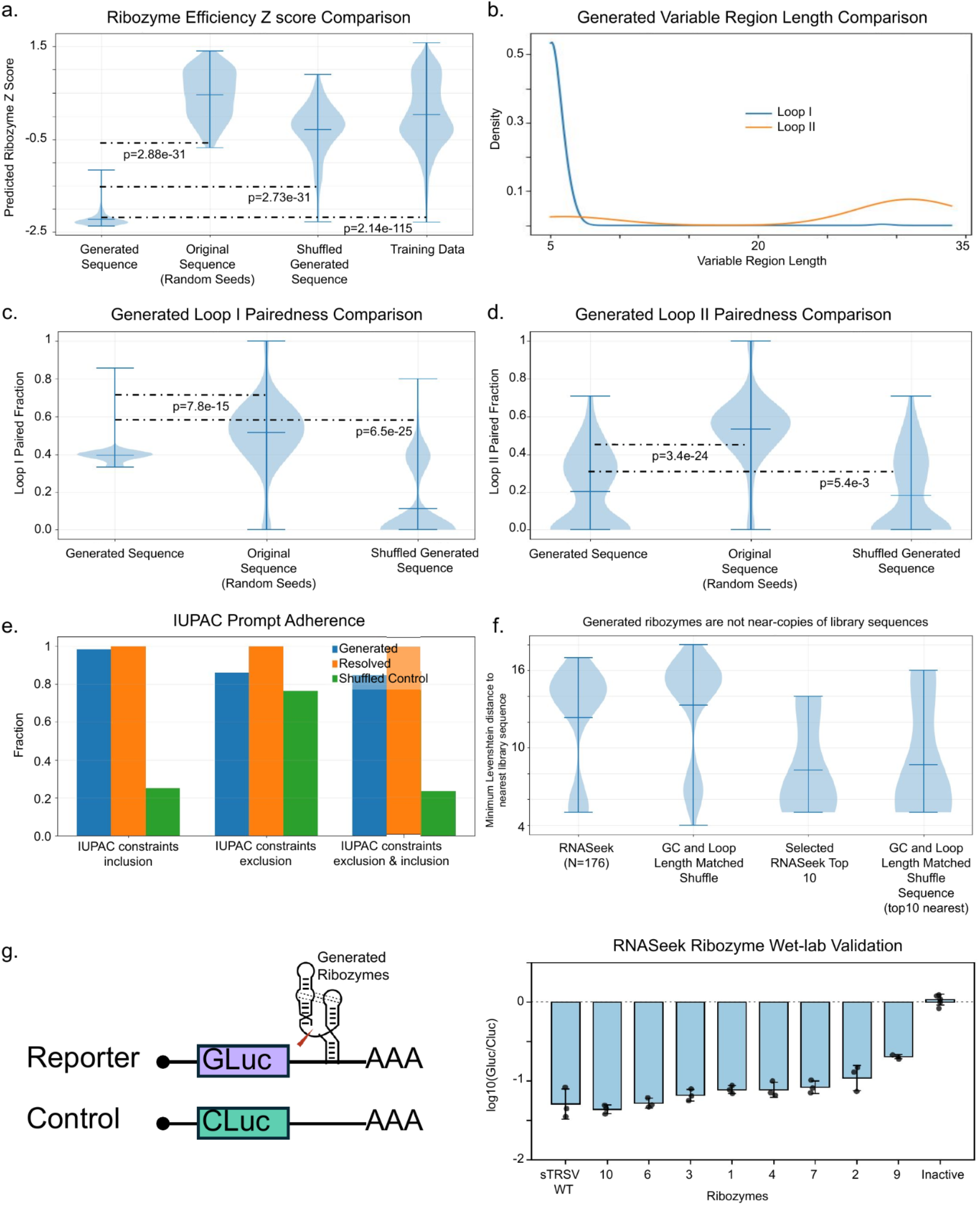
RNASeek generates active ribozymes through reward-guided sequence optimization. a. RNASeek was used to generate ribozyme sequences, which were evaluated by the trained reward model against IUPAC-resolved sequences, shuffled RNASeek outputs, and sequences from the wet-lab ribozyme library used for training. Lower predicted z-score values indicate higher ribozyme activity. b. Distribution of Loop I and Loop II lengths among RNASeek-generated ribozymes. c. Distribution of predicted pairedness within Loop I across RNASeek-generated ribozymes. d. Distribution of predicted pairedness within Loop II across RNASeek-generated ribozymes. e. RNASeek was used to synthesize ribozymes that include or exclude specific motifs through natural-language prompts. Instruction-following ability was benchmarked against shuffled controls and IUPAC-resolved sequences (Resolved), which were generated computationally to satisfy the IUPAC constraints without reward-model optimization. f. For each RNASeek-generated or control ribozyme sequence, the minimum Levenshtein distance to the closest sequence in the wet-lab ribozyme library was computed. Higher distances indicate lower similarity to known library sequences. RNASeek-generated sequences were compared with GC- and loop-length-matched shuffled controls, including matched controls for the selected top candidates. This analysis evaluates whether generated ribozymes are unusually close to wet-lab/library sequences, which would suggest memorization or near-copying of library sequences. g. Experimental validation of RNASeek-generated ribozymes using a dual-luciferase reporter assay, with GLuc under the control of the ribozyme in the 3′ UTR. In the bar plot, the x-axis represents different ribozymes, including wildtype sTRSV hammerhead ribozyme and an inactive ribozyme control, while the y-axis shows the log10-transformed GLuc/CLuc ratio normalized to the inactive ribozyme. Lower values indicate higher ribozyme activity.

### RNASeek discovers sequence and structural determinants of activity

Structural analysis of RL-generated ribozymes revealed several nontrivial design patterns. First, the two variable loops adopted distinct length distributions: Loop I clustered at short lengths while Loop II was broadly distributed in the 25–30 nt range (Figure 7B), recapitulating the short-loop/long-loop architecture associated with high activity (Figure 3A). Second, the loops differed in structural regime: Loop I occupied a narrow intermediate pairedness window, avoiding both the over-constrained profiles of the original sequences and insufficient pairing that would destabilize the local fold. By contrast, Loop II was structurally more permissive (Figures 7C–D). The shuffled controls disrupted both distributions, confirming that these structural signatures are sequence-encoded rather than compositional artifacts. Third, despite functional optimization, RNASeek maintained high adherence to prompt design constraints across motif inclusion, exclusion, and combined tasks (Figure 7E), with only modest reduction under the more complex simultaneous-constraint condition.

### RNASeek generates experimentally active ribozymes

We next assessed whether RNASeek-designed ribozymes represented novel sequence designs rather than near copies of sequences present in the wet-lab library. To evaluate this, we computed the minimum Levenshtein distance between each RNASeek-generated ribozyme and its nearest library sequence and compared these distances with those of GC- and loop-length-matched shuffled controls (Figure 7F). The full RNASeek set showed a distance distribution comparable to the matched shuffled sequences. Although the selected top RNASeek candidates had lower nearest-library distances, their GC- and loop-length-matched shuffled controls showed a similar distribution, indicating that this similarity is largely explained by sequence length, GC content, and loop constraints rather than direct sequence copying. These results provide no evidence that RNASeek simply reproduced or generated near copies of library sequences.

We then experimentally validated selected RNASeek-generated ribozymes using a dual-luciferase reporter assay, in which ribozyme-mediated cleavage reduces GLuc relative to the CLuc control (Figure 7G). All tested RNASeek-generated ribozymes substantially reduced reporter activity relative to the inactive control, demonstrating robust ribozyme activity in cells. Several generated ribozymes consistently approached the activity of the wild-type sTRSV ribozyme. Overall, the selected RNASeek-generated ribozymes consistently exhibited robust cleavage activity. Unlike 3′ UTR-mediated regulation of RNA stability, ribozyme self-cleavage is an intrinsic property of the RNA and therefore is expected to translate across reporter constructs and cellular contexts. Together, these results support that RL-guided RNASeek optimization can generate functional ribozyme designs while extrapolating beyond simple memorization of known wet-lab sequences.

## Discussion

### RNASeek as a cross-phyla RNA foundation model

In this study, we developed RNASeek, a multipurpose RNA language model capable of both predictive regression and sequence generation. RNASeek contains 1.6 billion parameters and was pretrained on a cross-phyla corpus spanning plant, animal, algal, and viral transcriptomes, together with natural-language descriptions and RNA structural information. Its unified input representation allows diverse tasks to be addressed using the same pretrained backbone, without requiring a separate model architecture for each application, and supports direct RNA sequence generation from natural-language-like instructions. Embedding analyses further show that RNASeek learns species-specific representations without explicit taxonomic labels, capturing organism-associated features such as motif usage, codon bias, and nucleotide composition through large-scale pretraining. RNASeek also demonstrates above-random zero-shot intron prediction, suggesting that these learned representations may support species-specific intron-related tasks without additional training.

A unified predictive and generative framework offers several advantages over pipelines in which sequence scoring and design are performed by separate architectures. Predictive performance provides an initial measure of whether the pretrained representation captures biologically relevant relationships within a given dataset and whether the available training data contain sufficient signal for the intended design objective. Although prediction accuracy does not by itself guarantee the quality of generated sequences, it provides an internally consistent basis for evaluating and optimizing candidates produced by the same pretrained backbone. Moreover, interpretation of the predictive model can identify sequence features associated with the target phenotype, such as motifs, nucleotide composition, positional effects, or structural properties. These analyses can provide hypotheses about the biological determinants of model behavior and help assess whether generated sequences achieve their predicted properties through plausible mechanisms rather than through obvious artifacts or unintended correlations.

The natural-language component further expands the utility of RNASeek by providing a flexible interface between biological objectives and sequence-level operations. Instead of defining every task through a new architecture or rigid numerical input format, users can describe desired sequence classes, organisms, structural features, functional constraints, or design objectives through prompt-based instructions. This capability could enable future applications such as conditional generation across species, simultaneous specification of multiple sequence constraints, interactive refinement of candidate RNAs, explanation or annotation of RNA features, and integration of experimental metadata or literature-derived knowledge into the design process. More broadly, natural-language conditioning may make RNA design models easier to adapt to new biological questions for which only limited task-specific data or computational infrastructure are available.

Recent RNA language models^6^ have shown that multi-transcriptome-trained transformer models can similarly capture evolutionary distance, codon bias, RNA-binding protein preferences, and high-level regulatory syntax across species. The RNA generative model that is closest in scale to RNASeek to date is GEMORNA^8^, which uses encoder–decoder modules to generate functional mRNAs with enhanced expression. However, GEMORNA remains smaller in scale (GEMORNA-CDS ∼4.22 million parameters; GEMORNA-5′ UTR ∼9.62 million parameters; GEMORNA-3′ UTR ∼37.28 million parameters) and does not combine long-context FlashAttention^38^-based training with modern reinforcement learning optimization such as GRPO. By contrast, RNASeek leverages FlashAttention for long nucleotide sequences, uses a specialized BPE tokenizer to encode UTRs, introns, and secondary structure with markup tokens, and demonstrates that an autoregressive decoder can support both de novo sequence generation and regression modeling within a unified architecture.

Accordingly, the benchmark results position RNASeek near the leading predictive models rather than as a uniformly superior predictor. The principal value of the framework is its unified objective: the same autoregressive backbone supports quantitative prediction, prompt-conditioned generation, and GRPO optimization, allowing functional scoring and sequence design to remain within a single model rather than requiring a specialized predictor to be coupled to a separate generator.

### Hammerhead ribozyme activity prediction, mechanistic rules, and generative design

In this section, we use hammerhead ribozymes as a model to study self-cleaving ribozymes and derive mechanistic rules. Predicting expression outcomes from this library has proven challenging for both RNASeek and traditional deep learning models. Nevertheless, RNASeek maintained higher prediction performance than shallower LSTM models, indicating that pretraining on a diverse transcriptomic corpus is beneficial for the model to develop a general understanding of how local sequence composition and long-range pairing tendencies combine to shape catalytic activity.

RNASeek organizes hammerhead ribozymes into discrete embedding clusters in intermediate model layers, reflecting distinct efficiency regimes and modes of catalytic action rather than treating the sequence space as a smooth, undifferentiated landscape. Across these clusters, high-efficiency ribozymes tend to have low GC content and increased unpaired probability in loop regions, suggesting that flexible, weakly paired loops are important for catalytic activity. Importantly, the residuals of the regression head remain broadly distributed across the embedding manifold rather than concentrating in specific regions, suggesting that the remaining prediction errors are unlikely to arise from a single underperforming layer or localized representational bottleneck.

The ribozyme regression head, built on the same RNASeek backbone, proved effective as a reward model for guiding sequence generation. Through GRPO-based reinforcement learning, the generative model was steered toward producing ribozyme sequences with predicted—and subsequently validated—activities that matched those observed in the experimental library. The model also recovered key sequence features underlying ribozyme activity using the reward model as the sole optimization signal. Because the model retains the ability to include or exclude specified motifs through near-natural-language instructions, we demonstrated that reward-driven generation and user-specified design control can be achieved simultaneously.

We also acknowledge several constraints in our current analysis and experimental setup. Our training dataset is limited to hammerhead ribozyme libraries with fixed backbones and only Loop I and Loop II randomized. The sequence activity relationships represented in this dataset may be influenced by library construction or cloning biases that preferentially include certain structural variants while under-sampling others. Although dimensionality reduction using UMAP revealed informative patterns, the method’s nonlinear embedding may obscure additional covarying features underlying the observed clusters. Future work will therefore benefit from targeted perturbation experiments guided by RNASeek interpretation, such as systematic sequence modifications within clusters to validate inferred sequence–activity relationships.

### RNASeek for stable 3′ UTR sequence prediction and design

In this section, we evaluate RNASeek as both a 3′ UTR stability predictor and a generative sequence design tool. Viruses rely heavily on regulatory RNA functions involved in transcription, translation, and RNA stabilization. Interpretation of the regression model’s learned representations indicates that AU-rich elements, GC content, positional nucleotide composition, and the minimum free energy (MFE) of predicted secondary structures all contribute to 3′ UTR-mediated RNA stability. Recent studies have begun combining machine learning with UTR stability prediction; however, most existing methods are non-generative and passively score fixed input sequences, leaving sequence optimization to random mutagenesis strategies that neither leverage pretrained genomic knowledge nor efficiently explore the sequence space. In contrast, RNASeek uses a single GPT-based backbone and shared tokenization system to both predict stability and generate optimized 3′ UTR sequences, unifying sequence scoring and design within the same pretrained framework while leveraging information learned from large public transcriptomic datasets^9–11,39^.

To actively optimize sequences rather than relying on random mutation, we used the stability prediction model as a reward model together with verifiable sequence-level constraints, including IUPAC motif inclusion/exclusion rules and length requirements specified through natural-language-style instructions. The resulting RNASeek-designed 3′ UTRs showed stability- and structure-related signatures consistent with targeted optimization. Compared with length- and GC-matched random sequences, natural human 3′ UTR sequences, and ChatGPT-5-generated sequences, RNASeek-designed sequences showed consistently higher predicted stability and a concentrated MFE range. This suggests that reinforcement learning steered the model toward a high-predicted-stability regime rather than simply optimizing a single compositional feature. When CpG content was held constant and AU syntax was ablated, predicted stability rapidly deteriorated, suggesting that reinforcement learning used AU-rich syntax to regulate predicted stability (Figure 6F). These characteristics are consistent with our current understanding that AU elements and low GC content contribute to mRNA stability, while further suggesting that targeted poly(A)-like tract placement may contribute to the optimized sequence phenotype.

Finally, we highlight several caveats regarding generalization and baseline comparisons. Viral elements were evaluated only in the context of a luciferase reporter, and although 3′ UTR function is not necessarily CDS-specific, reinforcement learning may amplify reward-model biases favoring sequences compatible with this upstream coding context. Additionally, the ChatGPT-5 benchmark does not exhaustively evaluate all possible decoding strategies for a general-purpose language model.

### Significance, limitations, and future directions

Taken together, our results establish RNASeek as a cross-phyla RNA foundation model that unifies sequence representation, regulatory prediction, and reward-driven generative design within a single autoregressive framework. Conceptually, this work demonstrates that large language model architectures and policy-optimization methods developed for natural language can be adapted to biological sequence design, enabling both quantitative functional prediction and constraint-guided generative optimization within a unified framework. A fundamental limitation of the current study is that RNASeek does not explicitly encode cell type, transcriptomic state, or other context-specific cellular features as inputs. Reward-model accuracy therefore represents a ceiling on the quality of RL-guided design. In the future, related pretraining and reinforcement-learning pipelines could be extended to proteins and other biological macromolecules, although such extensions would require task-specific validation. The cross-phyla generalization demonstrated here suggests that a single large-scale biological language model, trained on sufficiently diverse sequence corpora, may serve as a shared representational backbone for a wide range of prediction and design tasks, including enzyme activity prediction, UTR stability modeling, splicing, subcellular localization, and therapeutic protein or RNA design. Future advances in multimodal reward modeling, more phylogenetically diverse pretraining corpora, and systematic experimental validation pipelines will likely strengthen and extend the RNASeek framework, making model-guided RNA design increasingly accurate, generalizable, and biologically informative.

## Acknowledgments

We would like to appreciate members of the Xiang and Jia lab for providing helpful discussions. We would also like to thank Xuan Chen, Ada Yang and Karen Zhao for critical feedback on the manuscript. Computations were performed using the computer clusters and data storage resources of the HPCC, which were funded by grants from NSF (MRI-2215705, MRI-1429826) and NIH (1S10OD016290-01A1).

## Author contributions

S.C. and J.S.X. conceived the project. S.C., J.S.X., and Z.J. designed the research. S.C. developed the software. S.C. and J.S.X. performed the bioinformatic analyses. S.C., J.S.X., Z.J., W.V.L. and L.W. analyzed the data. S.C. and J.S.X. designed the wet lab experiments. J.S.X. and N.F. performed the experimental validation. S.C. and J.S.X. wrote and edited the manuscript. W.V.L., L.W. and Z.J. edited the manuscript. J.S.X. and Z.J. supervised the project.

## Competing interests

The authors declare no conflict of interest.

## Code Availability

The source code used for RNASeek training, evaluation, sequence generation and model weights are available at https://huggingface.co/JoyXiangLab/rnaseek-full.

## Supplementary Figures

**Supplementary Figure 1.**
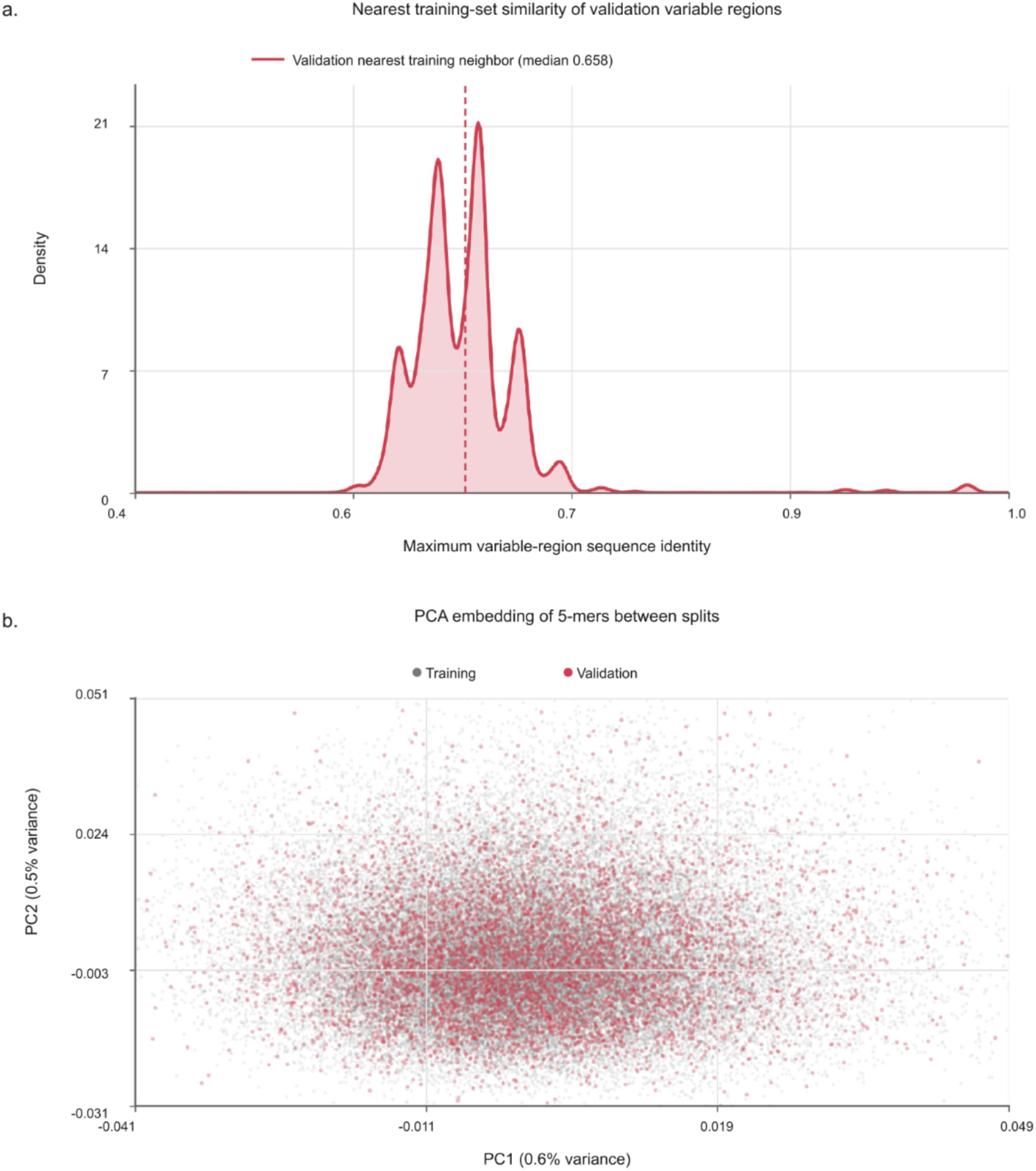
Train–validation design-space similarity. a. Density plot of nearest-neighbor sequence similarity between validation and training variable regions. Three invariant catalytic-core segments (GCTGTCACCGG, TCCGGTCTGATGAGTCC, GGACGAAACAGC) were removed from each sequence to isolate Loop I and Loop II. Each of the 5,756 validation examples was compared exhaustively with all 51,804 training examples. For each pair, Levenshtein distances were calculated separately for the two variable loops, summed, and normalized by their combined aligned lengths; the maximum identity for each validation example was retained. The dashed line marks the distribution median (0.658). The training and validation sets contain 0 exact variable-region overlaps. b. Principal component analysis of normalized 5-mer frequency embeddings calculated from Loop I and Loop II only, excluding 5-mers spanning the loop boundary. PCA was fitted on the 51,804 training examples, and the 5,756 validation examples were projected into the fitted space and overlaid. Training examples are shown in gray and validation examples in red. PC1 and PC2 explain 0.6% and 0.5% of training-set variance, respectively.

**Supplementary Figure 2.**
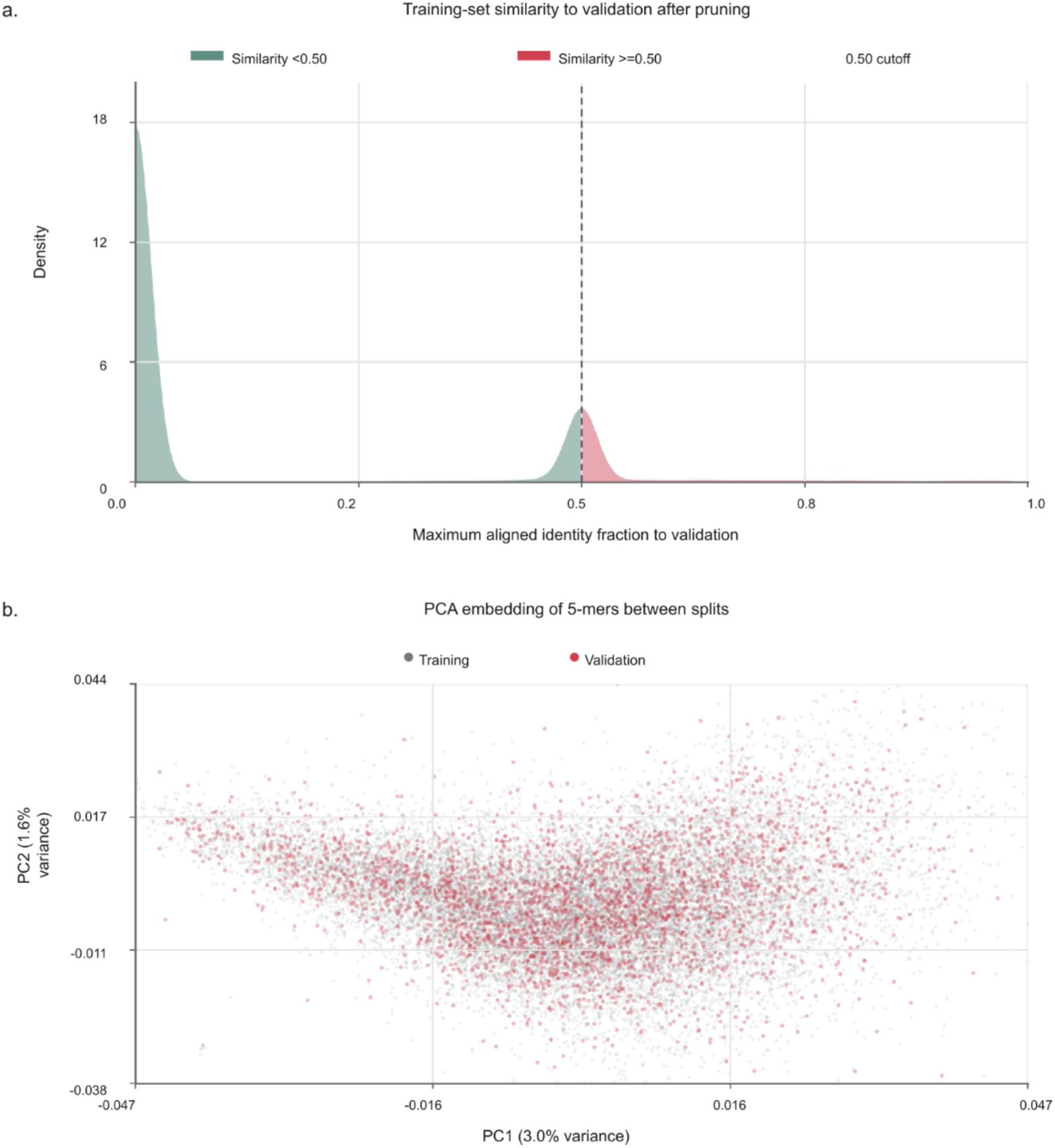
Train–validation sequence-space comparison after similarity filtering. a. Density of maximum local-alignment similarity from each original training viral insert to the validation set. Constant cloning flanks and the 7-nt barcode were removed before comparison, leaving the 130-nt viral insert. Similarity is shown as aligned identical nucleotides divided by the full insert length. Training sequences with an MMseqs2 forward-strand Smith–Waterman local nucleotide alignment satisfying at least 50% nucleotide identity and at least 50% query and target coverage to any validation insert were removed before model training (4,669 of 23,484 original training rows). A single smoothed density is colored on each side of the 0.50 threshold; the dashed line marks the cutoff. b. PCA embedding of normalized 5-mer frequency profiles calculated from the same 130-nt viral insert. PCA was fitted on the 18,815 retained training sequences, and the 2,935 validation sequences were projected into the fitted space and overlaid.

**Supplementary Figure 3.**
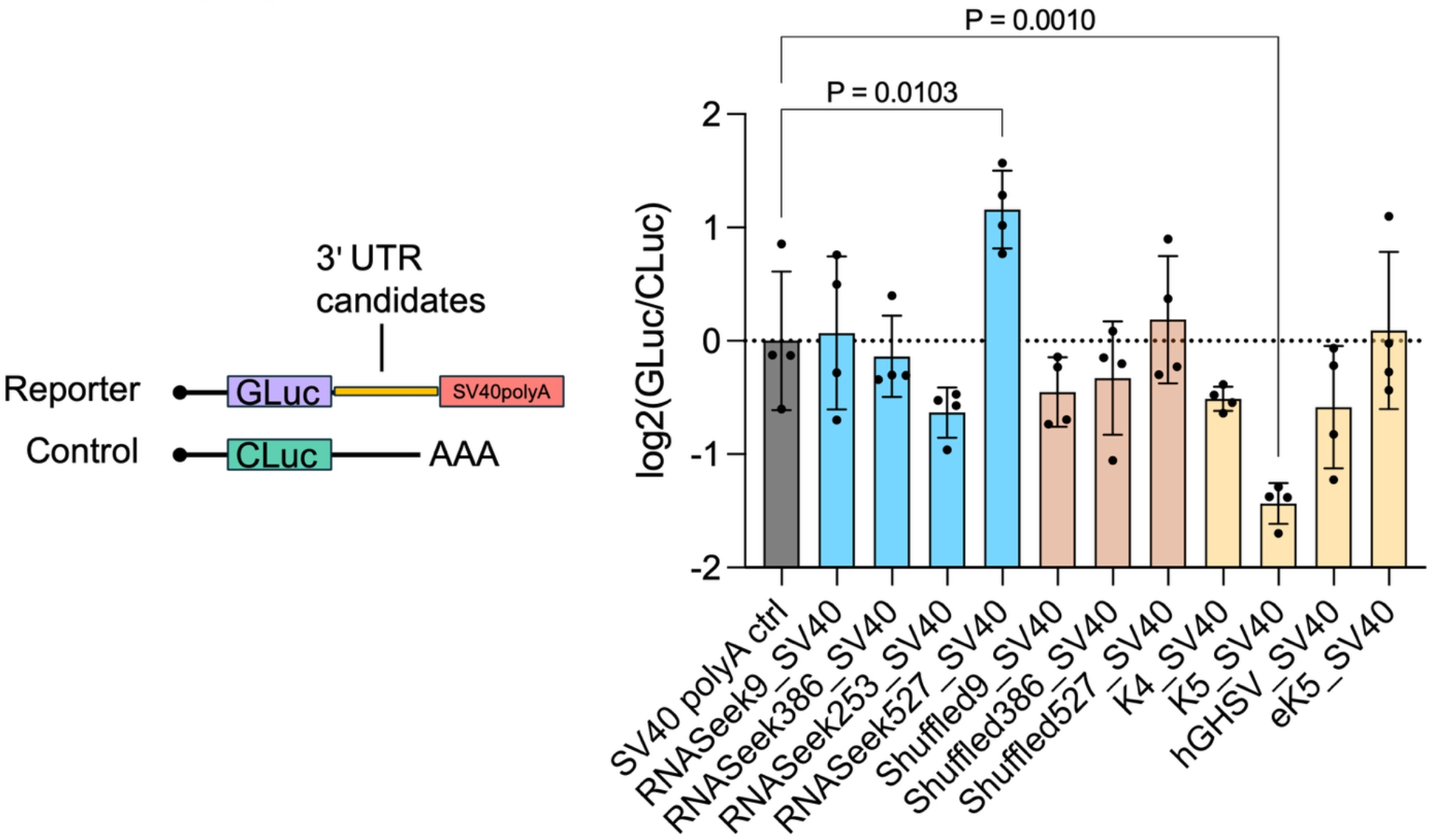
Regulatory activity in combination with the SV40 poly(A) sequence. Reporter activity of RNASeek-generated 3′ UTR candidates (blue), their corresponding shuffled sequences (copper), and benchmark sequences from the training data (yellow) was measured in the Gluc/Cluc reporter system in which the sequences were inserted between the stop codon and the SV40 poly(A) sequence. Bars show mean reporter activity, with individual data points representing biological replicates. Statistical significance was assessed using two-sided unpaired *t*-tests with Dunnett’s multiple-comparisons correction.

## Notes

https://huggingface.co/JoyXiangLab/rnaseek-full

### Competing Interest Statement

The authors have declared no competing interest.

## References

1. Wurmthaler, L. A., Klauser, B. & Hartig, J. S. Highly Motif- and Organism-Dependent Effects of Naturally Occurring Hammerhead Ribozyme Sequences on Gene Expression. (2018).

2. Zhou, J. & Troyanskaya, O. G. Predicting effects of noncoding variants with deep learning–based sequence model. Nature Methods 12, 931–934 (2015).

3. Alipanahi, B., Delong, A., Weirauch, M. T. & Frey, B. J. Predicting the sequence specificities of DNA- and RNA-binding proteins by deep learning. Nature Biotechnology 33, 831–838 (2015).

4. Jaganathan, K. et al. Predicting Splicing from Primary Sequence with Deep Learning. Cell 176, 535–548.e24 (2019).

5. Avsec, Ž., et al. Effective gene expression prediction from sequence by integrating long-range interactions. Nature Methods 18, 1196–1203 (2021).

6. Penić, R. J., Vlašić, T., Huber, R. G., Wan, Y. & Šikić, M. RiNALMo: general-purpose RNA language models can generalize well on structure prediction tasks. Nat Commun 16, 5671 (2025).

7. Ji, Y., Zhou, Z., Liu, H. & Davuluri, R. V. DNABERT: pre-trained Bidirectional Encoder Representations from Transformers model for DNA-language in genome. Bioinformatics 37, 2112–2120 (2021).

8. Zhang, H. et al. Deep generative models design mRNA sequences with enhanced translational capacity and stability. Science 10.1126/science.adr8470 (2025) doi:10.1126/science.adr8470.

9. Castillo-Hair, S. et al. Optimizing 5’UTRs for mRNA-delivered gene editing using deep learning. Nature Communications 15, 5284 (2024).

10. Pan, S. et al. UTR-Insight: integrating deep learning for efficient 5′ UTR discovery and design. BMC Genomics 26, 107 (2025).

11. Wang, L. et al. Machine learning-based analysis of the impact of 5′ untranslated region on protein expression. Nucleic Acids Res 53, gkaf861 (2025).

12. Chen, J., et al. Interpretable RNA Foundation Model from Unannotated Data for Highly Accurate RNA Structure and Function Predictions. (2022).

13. Wang, N. et al. Multi-purpose RNA language modelling with motif-aware pretraining and type-guided fine-tuning. Nature Machine Intelligence 6, 548–557 (2024).

14. Chen, K. et al. Self-supervised learning on millions of primary RNA sequences from 72 vertebrates improves sequence-based RNA splicing prediction. Brief Bioinform 25, (2024).

15. Dalla-Torre, H. et al. Nucleotide Transformer: building and evaluating robust foundation models for human genomics. Nature Methods 22, 287–297 (2024).

16. Nguyen, E., et al. HyenaDNA: Long-Range Genomic Sequence Modeling at Single Nucleotide Resolution. *ArXiv* (2023).

17. Lambert, N. Reinforcement Learning from Human Feedback. https://arxiv.org/html/2504.12501v3.

18. Eastman, P., Shi, J., Ramsundar, B. & Pande, V. S. Solving the RNA design problem with reinforcement learning. PLoS Comput Biol 14, e1006176 (2018).

19. Ahmad, J. M. et al. A curation system of rice trait ontology with reliable interoperation by LLM and PubAnnotation. Genomics Inform 23, 24 (2025).

20. Lambert, C. N. et al. Exploring the space of self-reproducing ribozymes using generative models. Nature communications 16, (2025).

21. Rotrattanadumrong, R. & Yokobayashi, Y. Experimental exploration of a ribozyme neutral network using evolutionary algorithm and deep learning. Nature communications 13, (2022).

22. Seo, J. J., Jung, S. J., Yang, J., Choi, D. E. & Kim, V. N. Functional viromic screens uncover regulatory RNA elements. Cell 186, (2023).

23. Xiang, J. S. et al. Massively parallel RNA device engineering in mammalian cells with RNA-Seq. Nature Communications 10, 4327 (2019).

24. Townshend, B., Kennedy, A. B., Xiang, J. S. & Smolke, C. D. High-throughput cellular RNA device engineering. Nature Methods 12, 989–994 (2015).

25. Strobel, B. et al. High-throughput identification of synthetic riboswitches by barcode-free amplicon-sequencing in human cells. Nature communications 11, (2020).

26. Townshend, B., Xiang, J. S., Manzanarez, G., Hayden, E. J. & Smolke, C. D. A multiplexed, automated evolution pipeline enables scalable discovery and characterization of biosensors. Nature communications 12, (2021).

27. De la Peña, M., Gago, S. & Flores, R. Peripheral regions of natural hammerhead ribozymes greatly increase their self-cleavage activity. EMBO J 22, 5561–5570 (2003).

28. Schmidt, C. M. & Smolke, C. D. A convolutional neural network for the prediction and forward design of ribozyme-based gene-control elements. eLife 10, (2021).

29. Martick, M. & Scott, W. G. Tertiary Contacts Distant from the Active Site Prime a Ribozyme for Catalysis. Cell 126, 309 (2006).

30. Dyer, S. C. et al. Ensembl 2025. Nucleic Acids Research 53, D948 (2024).

31. riboviria.org. https://riboviria.org.

32. Lorenz, R. et al. ViennaRNA Package 2.0. Algorithms for molecular biology : AMB 6, (2011).

33. Qwen et al. Qwen2.5 Technical Report. (2024).

34. Devlin, J., Chang, M.-W., Lee, K. & Toutanova, K. BERT: Pre-training of Deep Bidirectional Transformers for Language Understanding. (2018).

35. Bailey, T. L. STREME: accurate and versatile sequence motif discovery. *Bioinformatics (Oxford*, England) 37, (2021).

36. Cox, D. B. T. et al. RNA editing with CRISPR-Cas13. Science 358, 1019–1027 (2017).

37. Xiang, J. S. et al. Genome-Wide Interrogation of SARS-CoV-2 RNA-Protein Interactions Uncovers Hidden Regulatory Sites. bioRxiv (2025) doi:10.1101/2025.05.26.656146.

38. Dao, T., Fu, D. Y., Ermon, S., Rudra, A. & Ré, C. FlashAttention: Fast and Memory-Efficient Exact Attention with IO-Awareness. (2022).

39. Chaturvedi, M., Rashid, M. A. & Paliwal, K. K. Transformers in RNA structure prediction: A review. Comput Struct Biotechnol J 27, 1187–1203 (2025).

